# A multiscale chromatin collapse underlies vascular pathology in Hutchinson–Gilford progeria syndrome

**DOI:** 10.64898/2025.12.11.693713

**Authors:** Mzwanele Ngubo, Naveen Ahuja, Rana Karimpour, Amit Shrestha, Michael J. Hendzel, Theodore J. Perkins, William L. Stanford

## Abstract

Hutchinson-Gilford progeria syndrome (HGPS) is a devastating premature aging disorder driven by the accumulation of the lamin A variant, progerin, leading to severe vascular pathology. While epigenetic alterations are implicated, the spatiotemporal reorganization of the higher-order chromatin and its functional impact on vascular smooth muscle cell (VSMC) transcription remain poorly defined. Through an integrated, longitudinal multi-omics approach combining *in situ* high-throughput chromosome conformation capture (Hi-C) and Cleavage Under Targets and Tagmentation (CUT&Tag) profiling of CTCF, SMC1A, H3K27me3, H3K27ac, and H3K36me3 with transcriptomic analyses from control and HGPS iPSC-derived VSMCs, we reveal that global topologically associating domain (TAD) architecture remains largely intact in HGPS. However, the internal chromatin states of TADs undergo dynamic, passage-specific remodeling, characterized by a progressive accumulation of broad H3K27me3-repressed domains. This is accompanied by a loss of A/B compartment segregation, as confirmed by DNA-FISH, and a pronounced depletion of interchromosomal intermingling, specifically surrounding downregulated genes, suggesting that these interchromosomal contacts are essential for maintaining transcriptional competence in normal VSMCs. Crucially, we uncover widespread rewiring of enhancer-promoter (E-P) loops, which is linked to the dysregulation of genes critical for vascular development, extracellular matrix organization, and atherosclerosis. Our study demonstrates that spatiotemporal redistribution of repressive histone marks and reorganization of E-P interactions within a structurally resilient TAD framework underpin widespread transcriptional dysregulation in HGPS vascular pathogenesis. This uncovers a critical dissociation between higher-order chromatin architecture and histone modification landscape, providing a mechanistic basis for the failure of vascular homeostasis in progeria.

## 1 Introduction

Hutchinson-Gilford progeria syndrome (HGPS) is a rare, fatal premature aging disorder caused by a heterozygous *de novo* mutation in the *LMNA* gene, leading to the production of progerin, a truncated and toxic variant of lamin A [1,2]. Progerin disrupts nuclear architecture by accumulating at the nuclear periphery, destabilizing the nuclear lamina, and inducing hallmark cellular phenotypes such as genomic instability, nuclear deformities, loss of heterochromatin, and premature senescence [3,4]. These defects mirror aspects of physiological aging, positioning HGPS as an important model for studying aging-related chromatin and epigenetic dysregulation [4].

Central to HGPS pathology is the interaction between nuclear structure and epigenetic regulation. The nuclear lamina serves as a scaffold for chromatin organization, anchoring lamina-associated domains (LADs) and maintaining the spatial segregation of active (A) and inactive (B) chromatin compartments [5–7]. In healthy cells, these interactions ensure stable transmission of chromatin states through cell division, a process known as epigenetic inheritance, in which histone modifications, chromatin domains, and nuclear positioning are faithfully re-established after mitosis to maintain lineage identity and transcriptional fidelity [8,9]. However, in HGPS, progerin exerts dominant-negative effects that destabilize these lamina-chromatin interactions [3]. This instability manifests as loss of lamina-associated heterochromatin, redistribution of histone marks, and transcriptional noise that accumulate over time, features that parallel aging and interpreted as resulting from defective epigenetic inheritance. In vascular smooth muscle cells (VSMCs), where mechanical stress and high turnover demand precise epigenetic maintenance, such inheritance defects accelerate loss of contractile identity and promote pro-atherogenic transcriptional programs, contributing to vascular stiffening and atherosclerosis characteristic of both HGPS and age-related cardiovascular disease [10,11].

Multiple studies have reported widespread alterations in histone modifications and chromatin-modifying activities in HGPS, including reduced levels of heterochromatin-associated marks such as H3K9me3 and H3K27me3, alongside increased H4K20me3 [12–16]. The depletion of H3K27me3 has been linked to decreased expression of the polycomb repressive complex 2 (PRC2) enzymatic subunit EZH2. Importantly, redistribution of H3K27me3 to specific gene promoters correlates with transcriptional dysregulation in HGPS, supporting a causal relationship between altered histone methylation and disease-associated gene expression changes. Together, these findings support a model in which progerin-mediated lamina disruption destabilizes heterochromatin–lamina interactions, driving progressive epigenetic reprogramming that contributes to vascular pathology and premature aging.

These findings have positioned chromatin dysfunction as a central feature of the disease. Yet, the mechanisms by which chromatin regulation functions across various spatial scales—from higher-order chromatin architecture to enhancer-promoter interactions—and its role in the progression of vascular diseases remain unclear. To investigate the epigenetic and chromatin-centric mechanisms underlying HGPS-driven vascular aging, we utilized VSMCs differentiated from HGPS patient-derived induced pluripotent stem cells (iPSCs) via an established directed differentiation strategy [17,18]. This model recapitulates replicative aging *in vitro*, allowing us to track the onset and progression of chromatin dysfunction over time. Crucially, our analysis includes pre-senescent cells (Passage 7), where progerin is detectable in ∼30% of the VSMC population, providing a window into early disease mechanisms. By comparing this pre-senescent state to a late-stage culture containing senescent cells (Passage 14, >50% progerin-positive), we establish a temporal framework to dissect how progressive lamin A/C dysfunction drives vascular aging. By integrating high-resolution chromatin capture (Hi-C), chromatin architectural proteins, histone modification analysis and high-resolution microscopy, we provide an invaluable resource elucidating the impact of progerin on the epigenome, chromatin topology, and cardiovascular gene regulation, offering insights into both HGPS and broader aging-related epigenetic decline.

## 2 Results

### Characterization of the chromatin states of TADs

We generated a 5kb high-resolution chromatin interaction map using *in situ* Hi-C to quantitatively assess changes in hierarchical chromatin architecture during vascular aging in two cell lines of control and HGPS iPSC-derived VSMCs, sampled at early passage (P7), representing the onset of significant progerin accumulation [17,18], and passage 14 (P14), characterized by advanced *in vitro* replicative aging with >50% progerin-positive cells. We first evaluated genome-wide contact probability as a function of genomic distance for both control and HGPS VSMCs. The similar chromatin contact decay profiles observed across samples at both early (P7) and late (P14) passages demonstrate robust comparability in Hi-C data quality, confirming equivalent sequencing depth, mappability, and filtering efficacy (Fig. 1a). This technical consistency establishes a foundational benchmark for subsequent analyses of chromatin organization. We assessed TAD boundary conservation between control and HGPS VSMCs at both early and late passages (Fig. 1b). At P7, 74.83% of TAD boundaries were conserved between control and HGPS cells, indicating a high degree of structural similarity at early passage. This level of conservation was further maintained at P14, where 79.06% of TAD boundaries overlapped between the two conditions, suggesting that large-scale chromatin domain organization remains broadly stable despite disease phenotype progression and replicative stress.

**Fig. 1:**
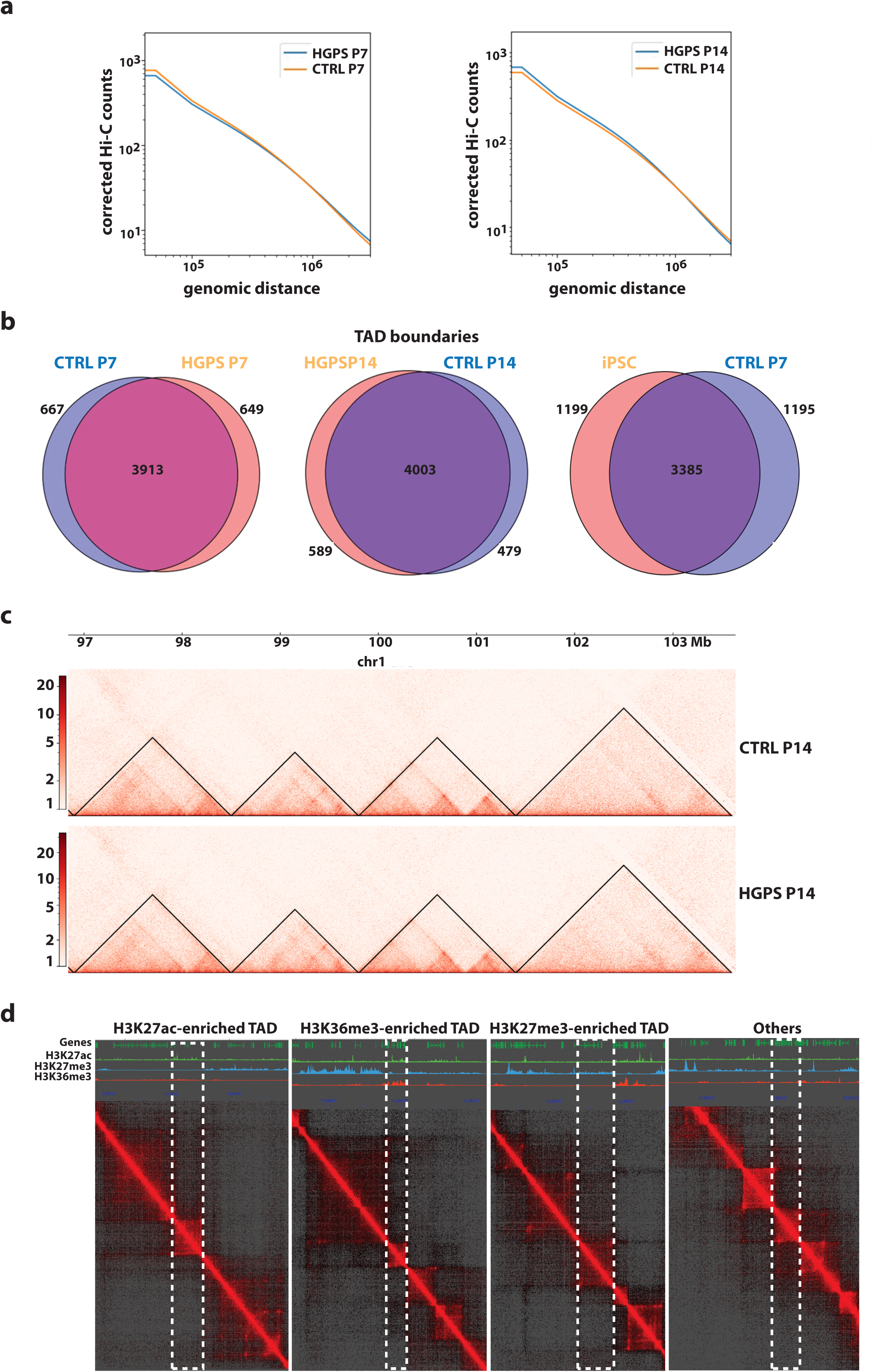
Overview of 3D genome organization. **a** Normalized Hi-C counts enrichment of control and HGPS VSMCs for both passage 7 and 14 across whole chromosomes at 50kb resolution. **b** Venn diagram depicting number of TAD boundaries overlapping and non-overlapping between control and HGPS samples for both passage 7 and 14, and between control P7 and iPSCs (4D nucleome project Accession: 4DNESPDEZNWX). **c** Comparison of Hi-C interaction map with annotated TADs between control and HGPS in chr1:96799888-103500000 for both passage 7 and 14 at 5kb resolution. Red indicates more frequent interactions and white indicates no interactions. **d** Hi-C contact maps showing TAD groups enriched in specific histone marks, with boundaries highlighted in white. Histone modification tracks include H3K27ac (green), H3K27me3 (blue), H3K26me3 (red) and other representing enrichment in none.

As an additional validation, we compared TAD boundaries between control VSMCs at P7 and undifferentiated iPSCs (4D nucleome project Accession: 4DNESPDEZNWX). In this case, we observed a lower conservation of 58.57%, consistent with the expected chromatin reorganization accompanying cellular differentiation. The contact maps from a representative genomic region in chromosome 1 show highly similar TAD structures between control and HGPS VSMCs (Fig. 1c). This observation is in line with previous reports showing that disruption of lamins does not perturb global TAD structure [19,20]. TAD boundaries are modulated by and enriched for the insulator protein CCCTC-binding factor (CTCF) and the cohesin complex [21–23]. We validated the integrity of TAD boundaries by performing (Cleavage Under Targets and Tagmentation) CUT&Tag of CTCF using biological triplicates of VSMCs (i.e., from three control and three HGPS iPSC lines). By plotting normalized coverage of CTCF across 2kb upstream and downstream of TAD boundaries as well as inside the TADs, we observed depletion within the TAD regions and enrichment of peaks at the boundaries as expected. This profile is highly similar between control and HGPS at passage 7 and passage 14 (Fig. S1a, b).

To assess the nature of the chromatin state of the TADs we performed further CUT&Tag experiments using biological triplicates of control and HGPS iPSC-derived VSMCs with antibodies that demarcate active and inactive genomic regions. We used H3K27ac to mark active enhancers and promoters [24,25], H3K36me3 to identify actively transcribed regions [26,27], and H3K27me3 to label repressed regions within TADs [28]. Examples of TADs enriched for H3K27ac, H3K36me3, and H3K27me3 are shown in (Fig. 1d). Examining the sizes of the TADs, we found that H3K27me3-enriched TADs were larger than H3K27ac and H3K36me3-enriched TADs in both control and HGPS VSMCs at passage 7 and passage 14 (Fig. S1c, d), consistent with the fact that this mark is known to form broad heterochromatic domains [29]. Conversely, H3K36me3-enriched TADs were significantly smaller (adjusted P < 0.05, Mann-Whitney-Wilcoxon test) and exhibited markedly higher gene density (Fig. S1e, f), in agreement with the role of H3K36me3 in demarcating transcriptionally active, gene-rich regions [29]. Additionally, H3K27ac-enriched TADs were also smaller than H3K27me3-enriched TADs, reflecting their focal association with active regulatory elements such as enhancers and promoters rather than extended domains [24,25]. Notably, the “Other TADs” were the largest TADs and had the lowest gene density. We postulate that these could be H3K9me3-enriched TADs correlating with our understanding of the mark being located at repetitive elements and low gene density areas, such as LADs [30].

### TAD epigenome structure regulates HGPS transcriptome

Having characterized the epigenetic states of all TADs, we next assessed how the epigenetic state may influence the expression level of the genes within the TADs. To assess the transcriptional output of TADs defined by chromatin state (based on histone modifications), we integrated our previously published RNA-seq data from control and HGPS VSMCs [18]. Consistent with the known association of H3K36me3 with active transcription, TADs enriched for this mark displayed higher average gene expression levels compared to other TAD subgroups at both early (P7) and late (P14) passages in control and HGPS cells (Fig. 2a, b). In contrast, H3K27me3-enriched TADs exhibit low gene expression levels (Fig. S1c, d), consistent with the known association of this mark with broad facultative heterochromatic domains [29].

**Fig. 2:**
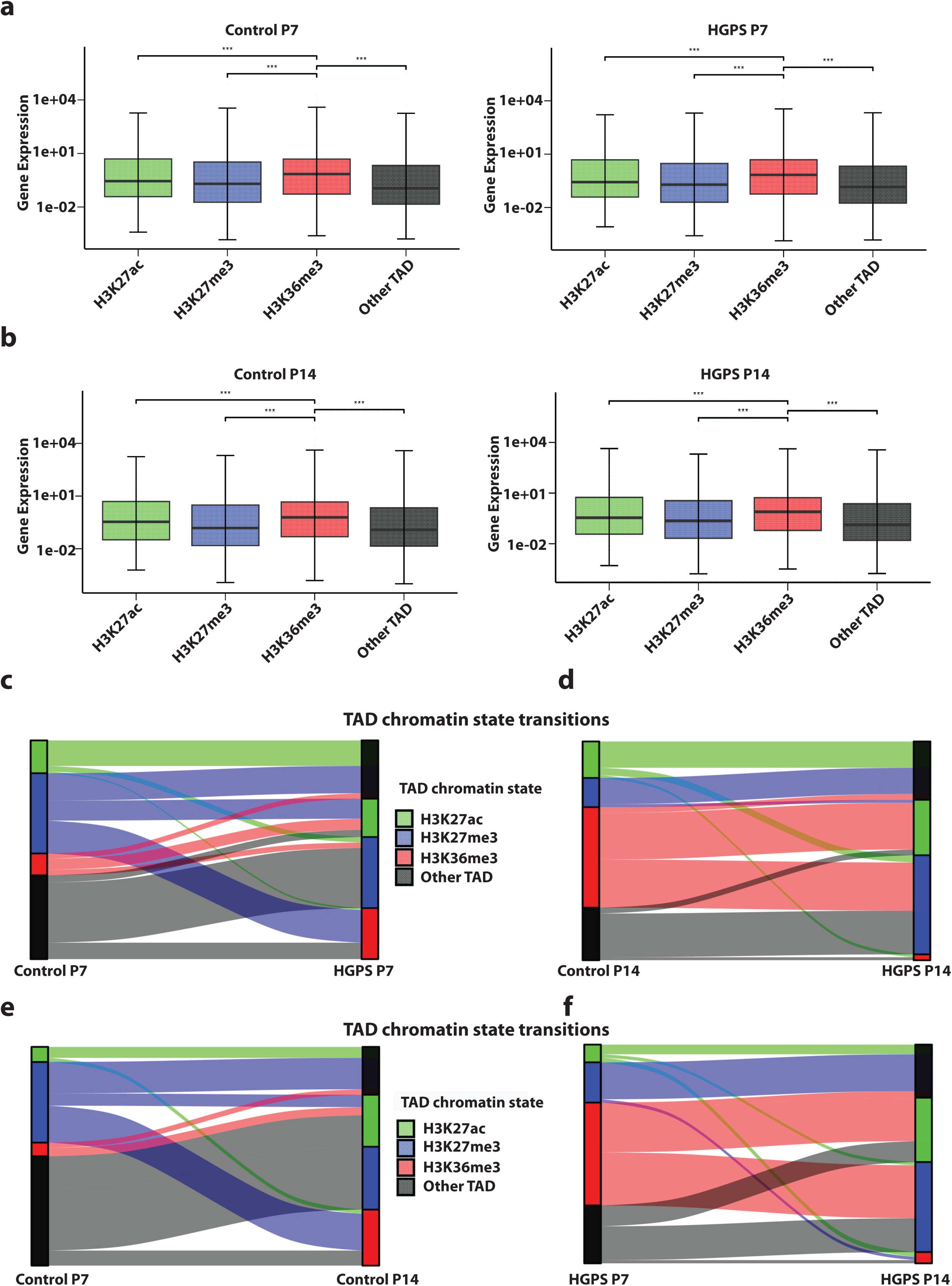
TAD structure regulates HGPS transcriptome. **a, b** Gene expression levels, measured in transcript per million (TPM) in histone mark enriched TADs. Panel **a** shows samples at passage 7, while panel **b** is at passage 14. Common TADs that changed epigenetic status between control and HGPS VSMCs. Panel **c** represents passage 7, while panel **d** is passage 14 of common TAD chromatin states transitions from control to HGPS VSMCs. Common TAD chromatin states transitions from control P7 to P14 (**e**) and HGPS P7 to P14 (**f**). P-values are computed using the Mann-Whitney-Wilcoxon test, *adj. p value < 0.05, **adj. p value < 0.01, ***adj. p value < 0.001.

To investigate how changes in TAD chromatin composition contribute to the transcriptional reprogramming observed in HGPS, we examined histone modification-defined chromatin states within TADs across passage 7 and 14. At passage 7, we identified 120 common TADs that switched chromatin state between control and HGPS cells (Fig. 2c and Table S1a-c). Of these, 18 TADs transitioned from H3K27me3 to H3K36me3, indicating a shift from a repressive to a transcriptionally active state. This transition is consistent with chromHMM analyses revealing the emergence of transcriptional activity within regions previously classified as heterochromatin (Fig. S2a). Additionally, 11 TADs switched from H3K27me3 to H3K27ac, representing a transition from facultative heterochromatin to an enhancer-associated active state. Conversely, 39 TADs gained H3K27me3 in HGPS, suggesting an alteration of repressive domains. By passage 14, the extent of chromatin remodeling expanded, with 150 common TADs undergoing chromatin state switches (Fig. 2d, Fig. S2b and Table S2a-c). Notably, 32 TADs transitioned from H3K36me3 to H3K27ac, indicating a switch from stable gene body transcription to enhancer-driven activation. This shift is corroborated by our chromHMM analysis, which detected the emergence of enhancer chromatin states within previously active transcriptional regions (Fig. S2b). Only two TADs switched from H3K27me3 to H3K27ac at this stage. Strikingly, 68 TADs gained H3K27me3 enrichment compared to 39 in passage 7, aligning with increased Polycomb repression chromatin states in chromHMM analysis. We also noted an overall increased peak width in HGPS VSMCs at passage 14 (Fig. S2c), reflective of spreading broad facultative heterochromatin during HGPS progression.

Next, we tracked TAD chromatin state transitions between passage 7 and 14 within each genotype. In control VSMCs, we detected 160 common TAD chromatin state transitions from passage 7 to 14 (Fig. 2e and Table S3). Of these, 46 TADs acquired H3K27me3, while 27 transitioned from H3K27me3 to H3K36me3, suggesting facultative heterochromatin remodeling. Additionally, 41 TADs gained H3K36me3, and 38 acquired H3K27ac, reflecting the coordinated activation of gene expression programs in normative aging. Whereas, HGPS VSMCs exhibited 136 TAD chromatin state switches over the same passages, but with a markedly different trajectory (Fig. 2f and Table S4). Specifically, 33 TADs switched from H3K36me3 to H3K27me3, and 27 transitioned from H3K36me3 to H3K27ac, indicating instability of transcriptionally active domains. The overall repression was more pronounced, with 56 TADs gaining H3K27me3, consistent with widespread Polycomb propagation. While 40 TADs acquired H3K27ac, these likely represent compensatory activation of regulatory elements amid global transcriptional dysregulation. Together, these findings reveal dynamic and stage-specific reorganization of chromatin states within TADs, with a predominant accumulation of H3K27me3-marked domains in HGPS. These structural chromatin alterations within TADs provide a spatial and molecular basis for the misregulation of gene expression characteristic of HGPS [31,32].

### HGPS shows loss of chromatin organization and transcriptional control at late passage

Given the chromatin state reorganization observed within TADs, we next asked whether these changes were accompanied by alterations in higher-order genome organization. To this end, we examined A/B compartment dynamics at passage 7 using dcHiC tool and identified 444 significant compartment changes between control and HGPS cells (Fig. 3a, Fig. S2d). This included 201 regions that transitioned from the transcriptionally active A compartment to the repressive B compartment, and 101 regions that shifted from B to A. Another category represents regions maintained their compartment status (76 A-to-A and 66 B-to-B) but showed significant changes in compartment strength as measured by principal component scores [33], reflecting altered chromatin activity despite preserved structural positioning, and indicating that early signs of compartmental reorganization are already detectable in HGPS. At passage 14, compartmental disruption increased, with 660 regions undergoing significant compartment changes (Fig. 3b, Fig. S2e). A total of 244 regions transitioned from A to B, while 160 regions switched from B to A. Although 124 and 132 regions retained A and B status, respectively, the overall increase in compartment switching at this stage suggests a remodeling of chromatin domain identity.

**Fig. 3:**
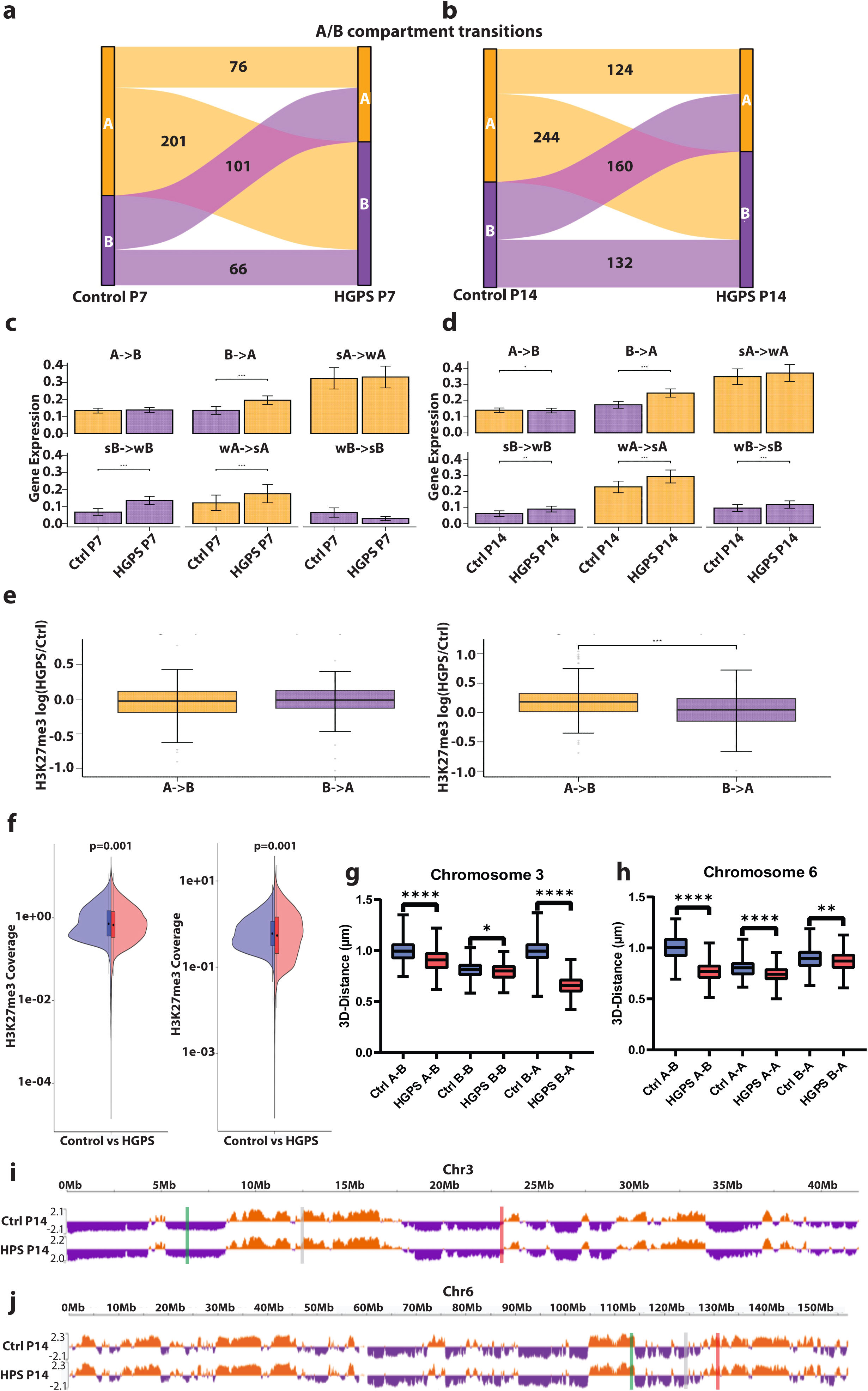
HGPS reveal loss of chromatin organization and transcriptional control at late passage. **a, b** Differential A/B compartment transitions at 50kb resolution (A compartments are represented in orange, while B compartments are in purple). Panel **a** shows samples at passage 7, while panel **b** is at passage 14. Gene expression at compartment transitions between control and HGPS. Strong (s) and weak (w) compartments were classified by comparing the relative compartment strength between samples. Panel **c** shows samples at passage 7, while panel **d** is at passage 14. **e** Comparison of H3K27me3 change at compartment transitions. The H3K27me3 log2 ratios (HGPS/control) are shown in the box plot for compartment transitions from A-to-B to B-to-A, across passage 7 (left panel) and passage 14 (right panel). **f** Differences in genome-wide B compartment H3K27me3 signal for control (blue) and HGPS (red) across passage 7 and 14. **g, h** Quantification of DNA-FISH 3D distances between A/B compartment assignments. Probes were designed along chromosomes 3 (**i**) and 6 (**j**) to alternate between regions belonging to the two compartments. P-values are computed using the Mann-Whitney-Wilcoxon test and Welch’s t test for compartment comparisons, *p value < 0.05, **p value < 0.01, ***p value < 0.001, ****p value < 0.0001.

To place these disease-associated changes in the context of normal cellular aging, we next analyzed compartment dynamics in control cells between passage 7 and passage 14. In contrast to HGPS, control aging was associated with a more limited degree of higher-order reorganization, with 135 significant differential A/B compartment switches detected over this interval (Fig. S2f). These included 40 A-to-B transitions and 40 B-to-A transitions, while 24 regions remained within the A compartment (A-to-A), reflecting subtle architectural modulation rather than wholesale compartmental reassignment.

Importantly, transcriptional consequences of compartment remodeling in aging control cells were modest (Fig. S2g). Both A-to-B and B-to-A transitions from P7 to P14 were associated with low gene expression, indicating that compartment switching during normal aging is largely uncoupled from immediate transcriptional output. Stratification by compartment strength further revealed no significant gene expression changes in regions transitioning from strong A to weak A or strong B to weak B, suggesting that weakening of compartment identity alone is insufficient to perturb transcription. In contrast, weak A-to-strong A transitions were associated with elevated gene expression, consistent with reinforcement of active chromatin states. Regions undergoing weak B-to-strong B transitions showed no significant transcriptional changes, indicating that heterochromatin consolidation during normal aging may be uncoupled from immediate transcriptional repression.

By comparison, compartment remodeling in HGPS cells exerted more pronounced and context-dependent effects on gene expression. Chromatin regions that have undergone A-to-B compartment switches did not exhibit significant changes in gene expression at early passage, suggesting that early-stage chromatin reorganization may precede transcriptional consequences (Fig. 3c). In contrast, B-to-A switches were associated with elevated gene expression, consistent with a gain in chromatin accessibility. No significant gene expression differences were observed in regions switching from strong A-to-weak A compartments. Strong B-to-weak B and weak A-to-strong A transitions correlated with increased gene expression. Whereas weak B-to-strong B regions showed a non-significant trend toward reduced expression, suggesting initial heterochromatin establishment. By late passage, however, compartment switching became more transcriptionally consequential as assessed by Wilcoxon Signed-Rank test (Fig. 3d). At P14, A-to-B transitions were accompanied by a reduction in gene expression, reflecting the repressive effect of late-stage heterochromatin expansion. Conversely, B-to-A switches continued to show elevated expression, consistent with reactivation of previously repressed regions. Strong A-to-weak A transitions did not yield significant changes in gene expression, suggesting retained transcriptional activity despite compartment weakening. In contrast, weak A-to-strong A transitions were associated with higher gene expression, consistent with reinforcement of active chromatin states. Similar to early passage, strong B-to-weak B transitions also exhibited increased gene expression. Surprisingly, in weak B-to-strong B transitions, typically associated with enhanced compartmental repression, we noted higher transcriptional output. These findings indicate that B compartment strengthening does not necessarily enforce gene silencing in late-passage HGPS cells.

We next determined whether compartment transitions in HGPS are accompanied by changes in repressive histone modifications. We examined H3K27me3 enrichment at regions undergoing A/B compartment switching. Comparing the log₂ ratio of H3K27me3 signal between HGPS and control VSMCs revealed distinct chromatin remodeling patterns associated with different compartmental shifts (Fig. 3e). Chromatin regions that switched from the active A compartment to the repressive B compartment did not exhibit significant differences in H3K27me3 enrichment compared to B-to-A regions in H3K27me3 signal between HGPS and control cells at passage 7. This suggests that early A/B compartment changes in HGPS are not yet coupled to repressive chromatin remodeling. Conversely, by late passage, A-to-B compartment switches relative to B-to-A regions displayed significantly higher H3K27me3 accumulation, indicating that loss of A compartment identity is increasingly accompanied by the spread of facultative heterochromatin. This is in agreement with a previous study in HGPS skin fibroblasts [34], where regions transitioning from open to closed compartments exhibited increased H3K27me3 enrichment and lamin A/C binding, while regions becoming more open showed a loss of both marks.

Building on the analysis of A/B compartment transitions, we sought to understand the relationship between chromatin compartmentalization and repressive epigenetic regulation in HGPS. We assessed global H3K27me3 levels within B compartments, which are typically associated with transcriptional repression and heterochromatic organization. Strikingly, B compartments in HGPS VSMCs exhibited a global reduction in H3K27me3 signal compared to controls at both early (P7) and late (P14) passages (Fig. 3f). This decrease was consistent across the genome-wide B compartments and was not restricted to specific compartment switch regions. The reduction in H3K27me3 within B compartments suggests a disruption of normal repressive chromatin architecture, despite the spatial maintenance of these domains. This is supported by findings in HGPS fibroblasts showing that regions with decreased H3K27me3 or lamin A/C binding often correspond to gene-poor, closed spatial compartments in normal cells, indicating a loss of chromatin repression even within structurally repressive domains [34].

We validated the A/B compartment assignments and directly assessed their spatial positioning within the nucleus by performing DNA fluorescence in situ hybridization (DNA-FISH) on chromosome 3 and 6 in control and late passage HGPS VSMCs. DNA-FISH enables direct measurement of 3D distances between compartment-assigned loci by fluorescently labeling specific A- and B-compartment probe pairs, allowing direct quantification of both homotypic (A–A, B–B) clustering and heterotypic (A–B, B–A) spacing. Thus, DNA-FISH provides a spatial readout of compartmental organization that can be compared to Hi-C–based assignments. In control cells, probe distances followed the expected compartmentalization pattern, with homotypic compartments positioned closer together than heterotypic compartments (Fig. 3g-i, Fig. S3a). For chromosome 3, B–B compartments were in closer spatial proximity (0.81 μm) compared to either B–A (0.99 μm) or A–B (1.00 μm) interactions, consistent with preferential clustering of repressed domains (Fig. 3g, i). For chromosome 6, A–A distances were shorter (0.80 μm) relative to both A–B (1.01 μm) and B–A (0.90 μm) interactions, reflecting the spatial coalescence of transcriptionally active regions (Fig. 3h, j). These control measurements thus establish that compartmental proximity is organized in a compartment-type–dependent and chromosome-specific manner. In late passage HGPS cells, these control-defined relationships were disrupted. On chromosome 3, the most striking change was a collapse of B–A segregation, with distances reduced from 0.99 μm in control to 0.66 μm in HGPS cells, indicating spatial mixing of inactive and active regions. B–B and A–B distances were only modestly affected (0.80 to 0.81 μm and 1.00 to 0.90 μm, respectively), indicating that loss of compartmentalization is driven primarily by erosion of heterotypic boundaries (Fig. 3g, i). On chromosome 6, A–B distances decreased from 1.01 μm in control to 0.77 μm in HGPS cells, again reflecting diminished segregation (Fig. 3h, j). In addition, A–A domains, which were tightly clustered in controls, became even closer in HGPS (0.80 to 0.74 μm), suggesting reinforcement of active-domain proximity despite increased A–B mixing. Together, these data reveal that while control VSMCs maintain compartment-specific organization, late passage HGPS VSMCs exhibit a loss of segregation between A and B domains, consistent with weakening of spatial compartmentalization in HGPS fibroblasts [34,35]. The observed architectural changes, specifically the preferential collapse of the inter-compartment distances, provide a direct structural explanation for the widespread transcriptional dysregulation in HGPS.

### Altered interchromosomal intermingling in HGPS VSMCs

To assess how interchromosomal intermingling changes with aging and premature aging, we analyzed Hi-C maps from control and HGPS VSMCs. We observed marked reduction in interchromosomal intermingling in HGPS compared to controls (Fig. 4a). To further characterize the cell-state-specific patterns, we generated difference maps derived from binarized Hi-C data that highlights regions preferentially engaged in interchromosomal contacts in each condition. Globally, 54.4% of intermingling regions were specific to control VSMCs, whereas only 31.4% were specific to HGPS while 14.1% were shared between control and HGPS, consistent with a pronounced depletion of interchromosomal interactions in HGPS cells (Fig. 4b).

**Fig. 4:**
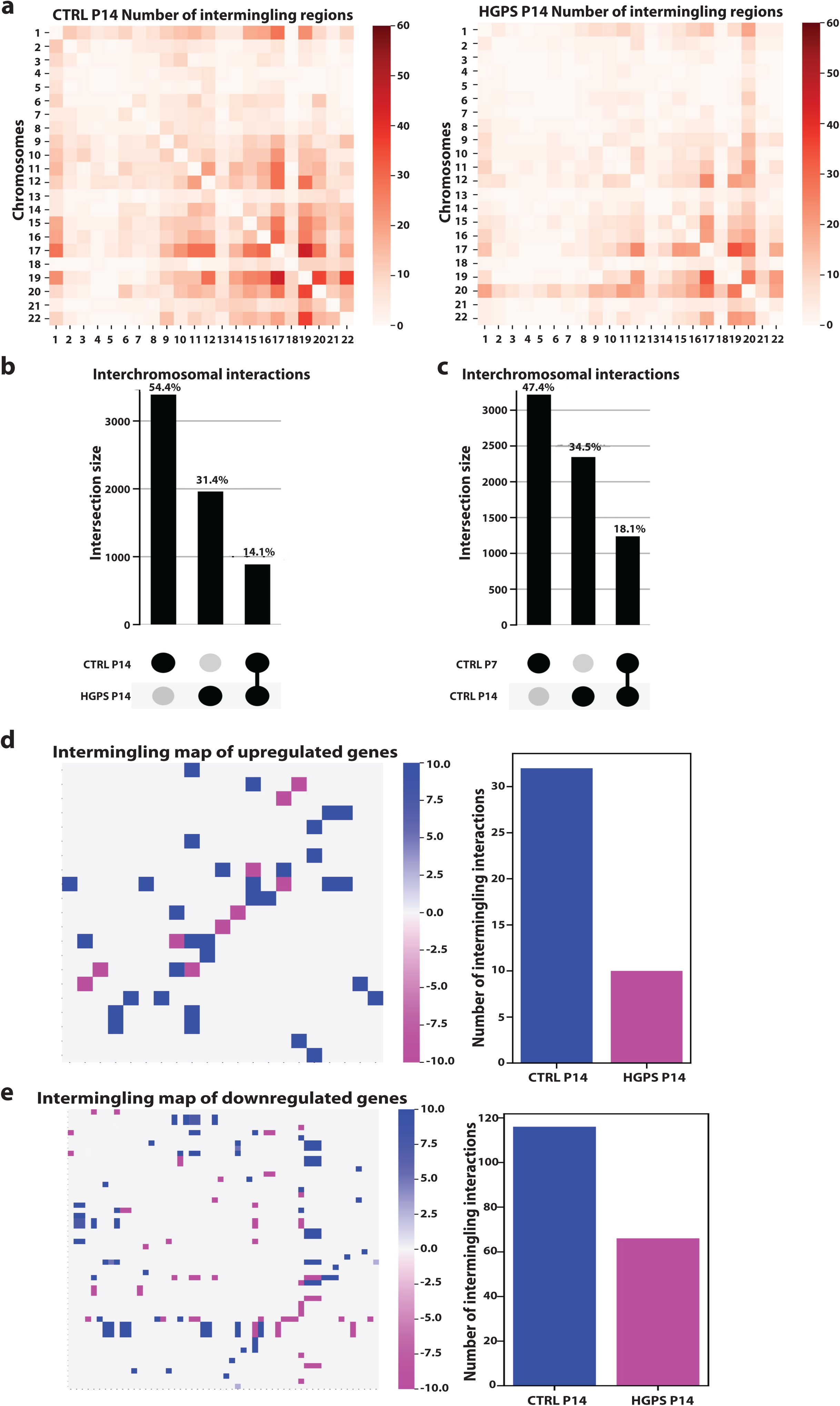
Reduced interchromosomal intermingling dynamics in HGPS VSMCs. **a** Genome-wide distribution and frequency of intermingling regions in control (left) and HGPS (right) samples, binned at a 250 kb resolution. **b** Interchromosomal large average submatrix (LAS) regions UpSet plot showing unique and overlapping 3D interchromosomal interactions between control and HGPS P14 VSMCs, and between control P7 and P14 VSMCs (**c**). **d**, **e** Intermingling difference map filtered for differentially expressed upregulated and downregulated genes in HGPS, positive values indicate interactions specific to control, negative values indicate interactions specific to HGPS, and zero represents shared interactions (left plot). Bar plot of total control and HGPS specific intermingling interactions in HGPS upregulated genes (right plot).

Changes in interchromosomal intermingling across normative aging were analyzed by comparing control P7 and P14 VSMCs (Fig. 4c). This analysis revealed a similar, though less pronounced, decline in interchromosomal intermingling, with 47.4% of regions specific to control P7, 34.5% specific to control P14, and 18.1% shared, indicating that loss of intermingling is a feature of physiological aging that is exacerbated in HGPS.

To understand how this relates to gene regulation, we analyzed intermingling patterns for upregulated genes in HGPS VSMCs. In control cells, these genes showed extensive interchromosomal contacts; however, in HGPS cells, these interactions were markedly reduced. This suggests that genes which become activated in HGPS are less engaged in long-range interchromosomal interactions. (Fig. 4d). Similarly, genes that are downregulated in HGPS also exhibited decreased interchromosomal intermingling. While these genes displayed higher levels of interaction in control cells, their interchromosomal contacts were notably diminished in HGPS. (Fig. 4e). Together, these results show that both upregulated and downregulated genes in HGPS are associated with reduced interchromosomal interactions, supporting a model in which premature aging leads to a global loss of genome intermingling that may contribute to widespread gene expression changes.

### Rewired enhancer–promoter chromatin loops are linked to transcriptional dysregulation of genes associated with vascular pathology in HGPS

To explore how alterations in enhancer–promoter (E–P) chromatin architecture contribute to transcriptional dysregulation in HGPS VSMCs, we examined genome-wide E–P loops and corresponding gene expression changes at early and late stages of disease progression. At the global level, genes associated with E–P loops showed a consistent trend toward higher expression in HGPS relative to control cells at both passages, reflecting elevated broad enhancer-mediated transcriptional activation in the disease context (Fig. 5a). To refine this analysis, we focused on differential E–P loops, those present in HGPS but absent in control, and vice versa. At passage 7, genes associated with differential E–P loops showed significantly increased gene expression in HGPS, indicating that early architectural rewiring is associated with altered transcriptional regulation (Fig. 5b). By passage 14, differential loops continued to be associated with a trend toward higher gene expression in HGPS, although the difference was not statistically significant, potentially reflecting progressive disorganization of regulatory chromatin and increased transcriptional variability.

**Fig. 5:**
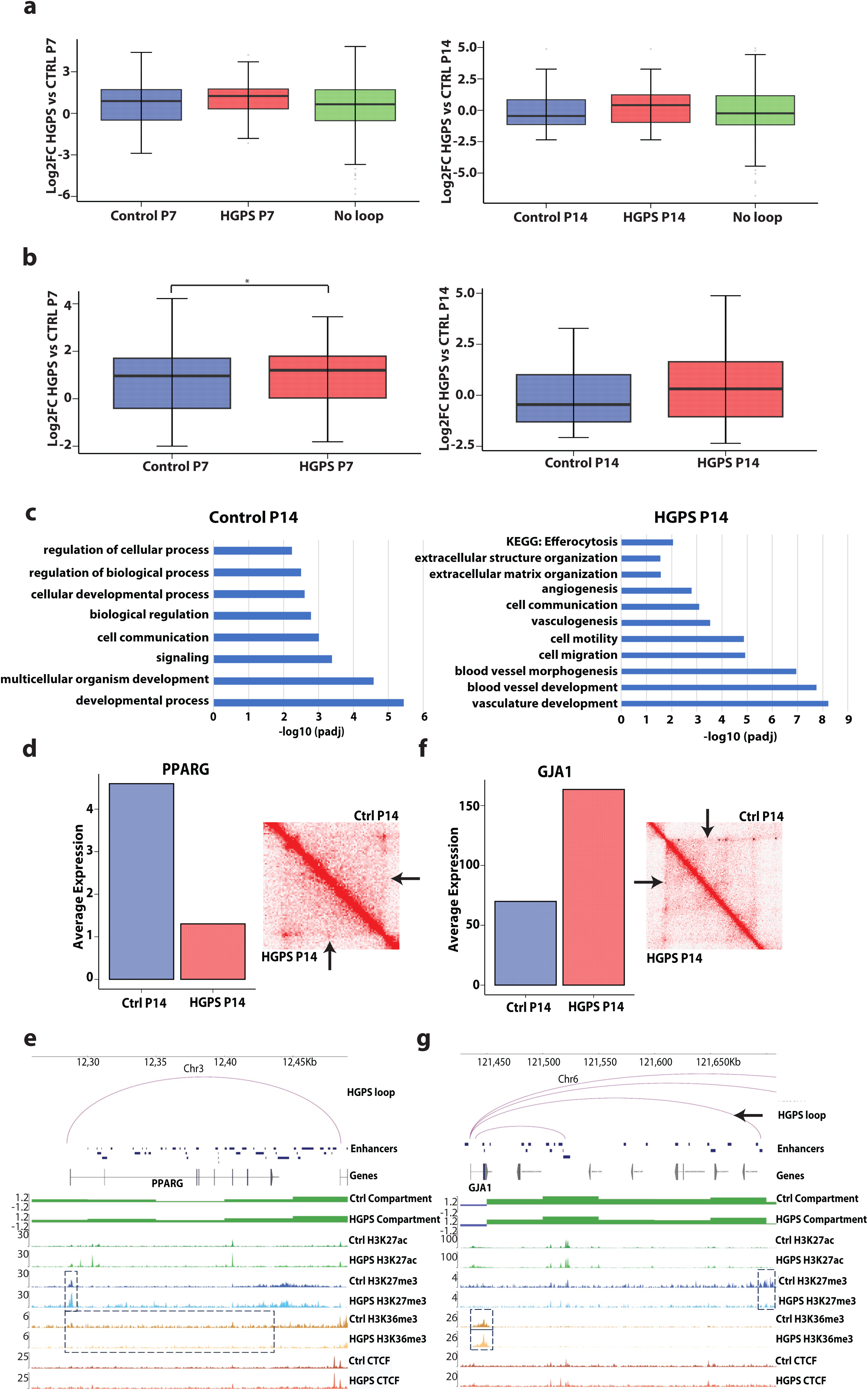
Differential enhancer-promoter chromatin loops promote dysregulation of genes associated with vascular pathology in HGPS. **a** Gene expression fold change between HGPS and control VSMCs, shown for genes with enhancer-promoter loops in control cells (blue), HGPS cells (red), or lacking loops (green). **b** Gene expression fold change (HGPS vs. control) associated with differential enhancer-promoter loops, those gained or lost in HGPS. **c** Gene ontology and KEGG pathway enrichment analysis for differential enhancer-promoter looped genes between control and HGPS at passage 14. Average expression (TPM) of PPARG (**d**) and GJA1 (left panel) (**f**). Hi-C interaction map of control and HGPS VSMCs at PPARG and GJA1 locus at 5kb resolution. The color intensity represents interaction frequency, while arrows indicate the PPARG and GJA1 differential enhancer–promoter loop (right panel). IGV genome browser showing enhancer–promoter loops, A/B compartments, CUT&Tag profiles of H3K27ac, H3K27me3, CTCF and SMC1A of PPARG (**e**) and GJA1 (**g**). P-values are computed using the Mann-Whitney-Wilcoxon test, *p value < 0.05, **p value < 0.01, ***p value < 0.001.

We next assessed the occupancy of architectural proteins at global loop anchors and observed consistently higher levels of CTCF signal overlapping with SMC1A, a key pair involved in chromatin loop formation and stabilization [36,37], in HGPS cells compared to controls at both passages (Fig. S3b, c), consistent with the increased TAD boundary strength observed in HGPS fibroblasts [35]. This suggests enhanced or aberrant loop formation in HGPS, potentially contributing to sustained enhancer–promoter interactions and the dysregulation of genes involved in vascular homeostasis and pathology.

Unbiased gene ontology and pathway enrichment analyses of differential E–P loop-associated genes at P14 identified biological processes driving HGPS vascular pathogenesis. In HGPS cells, differential E–P loops were significantly enriched for genes involved in vascular-related biological processes, including vasculature development, blood vessel morphogenesis (Fig. 5c). We further detected upregulated extracellular matrix production processes (extracellular structure organization and extracellular matrix organization), established drivers of atherosclerosis [38,39]. Genes involved in angiogenesis were also prominently associated with rewired E–P loops in HGPS cells. Aberrant angiogenic signaling within the blood vessel wall is a hallmark of advanced atherosclerotic lesions and contributes to intraplaque hemorrhage and increased risk of plaque rupture, both of which are clinically relevant complications in HGPS patients [40]. These processes are tightly linked to vascular remodeling and are consistent with pathological features observed in HGPS and atherosclerosis [41,42]. Moreover, KEGG pathway analysis uniquely identified efferocytosis, a macrophage-mediated clearance mechanism critical in the resolution of inflammation and atherosclerotic plaque remodeling [43], as enriched among genes associated with differential loops in HGPS VSMCs. This points to a potential functional link between enhancer–promoter rewiring and impaired clearance of apoptotic cells in diseased vascular tissue. In contrast, differential E–P loops in control cells were enriched for broader, homeostatic biological processes, such as regulation of cellular processes, developmental processes, cell communication, biological regulation, and signaling pathways. These categories reflect a more balanced and developmentally regulated transcriptional landscape, in sharp contrast to the vascular remodeling and stress-associated programs activated in HGPS.

These genome-wide patterns were exemplified by specific cases of disease-relevant regulatory rewiring. Notably, at passage 14, HGPS VSMCs exhibited a disease-specific enhancer–promoter loop at the *PPARG* locus, absent in control cells (Fig. 5d, Fig. S3d). Despite this physical interaction, *PPARG* expression was significantly reduced in HGPS, as confirmed by RNA-seq and qPCR (Fig. 5d, Fig. S4a). Chromatin profiling revealed elevated H3K27me3 at the promoter in HGPS (Fig. 5e). Additionally, reduced H3K36me3 over the gene body was noted in HGPS compared to control, suggesting transcriptional silencing despite structural promoter–enhancer connectivity. PPARG was reported to be downregulated in HGPS fibroblasts [44]. Studies have highlighted PPARG in VSMCs as a key determinant of atherosclerosis progression [45–47]. By maintaining VSMCs in a contractile, differentiated state and preventing their transition into a synthetic, pro-inflammatory phenotype, PPARG serves as a critical regulator of vascular integrity. Activation of PPARG in VSMCs reduces expression of matrix metalloproteinase-9 (MMP-9), which VSMCs and macrophages use, contributing to VSMC migration and plaque destabilization [48]. Its repression in HGPS, despite the presence of a regulatory loop, highlights a decoupling between chromatin architecture and transcriptional activation, likely driven by persistent facultative heterochromatin at regulatory elements.

In contrast, *GJA1* exhibited loop-associated activation rather than silencing (Fig. 5f). HGPS VSMCs displayed a disease-specific E–P loop at the *GJA1* locus, which was absent in control cells. *GJA1* expression was significantly upregulated in HGPS, as confirmed by both RNA-seq and qPCR (Fig. 5f, Fig. S4a). This activation coincided with increased H3K36me3 across the gene body in HGPS, consistent with active transcription, while control cells exhibited elevated H3K27me3 at the *GJA1* enhancer and gene body, indicative of gene repression (Fig. 5g). *GJA1*, a gap junction protein, promotes VSMC proliferation, migration, and phenotypic switching in atherosclerosis [49,50], and facilitates intercellular inflammatory signaling via calcium flux, ROS, and cytokines [51,52]. Its derepression in HGPS, linked to altered enhancer–promoter connectivity and chromatin remodeling, underscores how E–P loop rewiring can activate pathogenic vascular programs and contribute to disease progression. Additionally, we validated the differential enhancer-promoter loop interactions at PPARG and GJA1 by measuring pairwise 3D distances between enhancers and promoters using DNA-FISH probes at the PPARG and GJA1 loci. E–P distances were significantly shorter in HGPS than in control cells at both PPARG and GJA1 (Fig. S4b, Fig. S3d), consistent with the observed Hi-C chromatin looping. To assess loop specificity, we compared E–P distances to those between the enhancer and the midpoint of the loop. For both loci, enhancer–midpoint distances were consistently greater than E–P distances in both control and HGPS cells. Notably, enhancer–midpoint and promoter-midpoint distances did not differ significantly between conditions, confirming that the observed shortening was specific to E–P contacts rather than global chromatin collapse.

### HGPS aberrant gene expression coincide with altered epigenome

These gene-specific examples underscore how rewired enhancer–promoter interactions in HGPS can drive either aberrant activation or repression of vascular pathology genes, depending on local chromatin context. We next profiled the distribution of histone modifications associated with transcriptional repression (H3K27me3), elongation (H3K36me3), and activation (H3K27ac) at early and late disease stages. At passage 7, no significant differences in global H3K27me3 or H3K36me3 distribution were observed between HGPS and control cells (Fig. 6a, left panel; Fig. S4c, left panel). By passage 14, however, HGPS VSMCs exhibited a marked increase in H3K27me3 at gene promoters, consistent with the emergence of H3K27me3-enriched TADs (Fig. 6a, right panel; Fig. 2d; Fig. S2b and Table S2a-c). This local gain contrasted with a global reduction of H3K27me3 and EZH2 levels in HGPS and the loss of polycomb group bodies [53]. Whereas the control cells show multiple large distinct clusters of EZH2, consistent with polycomb group bodies, these are lost in HGPS cells. The redistribution of facultative heterochromatin was further reflected in reduced H3K27me3 occupancy at distal intergenic and intronic regions. Heatmaps of signals spanning 3 kb upstream of transcription start sites (TSS) to 3 kb downstream of transcription end sites showed increased H3K27me3 signal at TSSs in HGPS at passage 14 (Fig. S4d, e). Despite this shift, gene expression at newly marked H3K27me3 regions showed no significant change (Fig. 6c), and 12% of these regions also harbored H3K36me3 (Fig. 6d), suggesting that a subset of these regions undergo competitive crosstalk. In parallel, H3K36me3 showed no global changes at passage 7, but by passage 14, HGPS cells displayed a reduction in H3K36me3 signal within intronic regions and a modest increase at 5′ and 3’ UTRs (Fig. S4c), pointing to a reorganization of transcription elongation-associated marks. H3K27ac profiling revealed a global increase at promoters in HGPS, consistent with the formation of H3K27ac-enriched TADs (Fig. 6e; Fig. 2d; Fig. S2b and Table S2a-c), and a concurrent loss of H3K27ac in intronic regions, suggesting a coordinated reorganization of active and repressive chromatin regions. While promoter-proximal H3K27me3/H3K27ac shifts were topologically conserved, the dissociation between H3K27me3 accumulation and transcriptional silencing implies progressive gene regulatory failure.

**Fig. 6:**
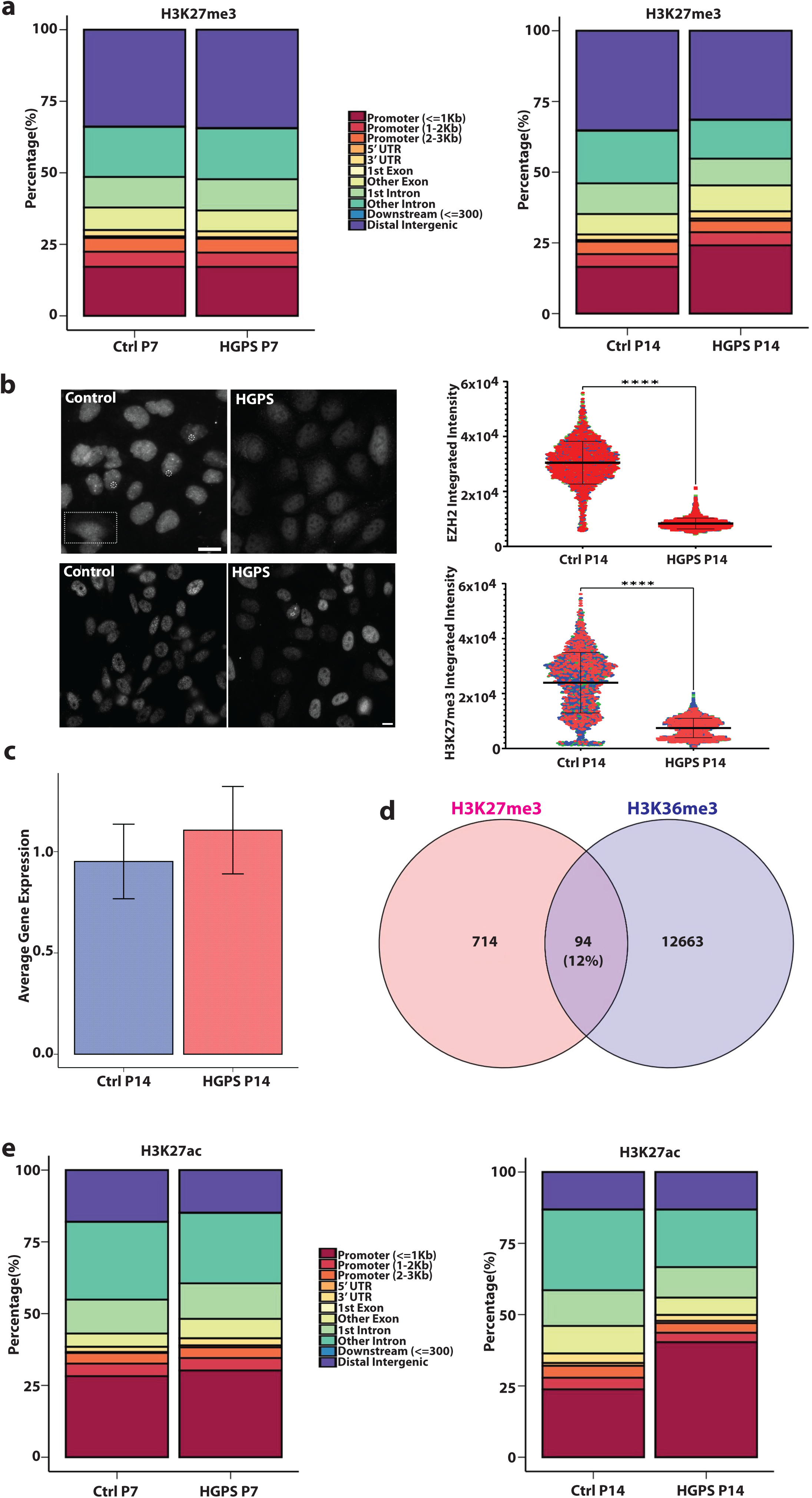
HGPS aberrant gene expression coincide with altered epigenome. **a** Genomic distribution of H3K27me3 peaks for control and HGPS at P7 (left panel) and P14 (right panel). **b** Immunofluorescence staining of VSMCs for EZH2 (top) and H3K27me3 (bottom). Images were acquired using identical exposure and scaling settings to enable quantitative comparison. Control cells display distinct nuclear EZH2 condensates (dashed circles), which are absent in HGPS VSMCs. A rare control cell shows cytoplasmic EZH2 with reduced nuclear signal (dashed rectangle), whereas this phenotype is common in HGPS cells, which overall exhibit reduced EZH2 levels and frequent cytoplasmic localization. HGPS VSMCs also show a global reduction of H3K27me3. **c** Average gene expression levels for genes enriched with redistributed H3K27me3 in HGPS compared to control at P14. **d** Venn diagram examining the overlap between H3K36me3 sites and newly gained H3K27me3 sites in HGPS P14. **e** Genomic distribution of and H3K27ac peaks for control and HGPS at P7 (left panel) and P14 (right panel). P-values are computed using the Mann-Whitney-Wilcoxon test, ****adj. p value < 0.0001.

## 3 Discussion

This study demonstrates that the chromatin architecture and epigenetic state are dynamically remodeled during vascular aging in HGPS, revealing a multilayered disruption of chromatin topology, histone modification patterns, and transcriptional regulation. By employing patient-derived iPSC–VSMCs and integrating high resolution Hi-C, histone modification profiling, and enhancer–promoter loop analysis across distinct aging passages, we establish a temporal framework linking chromatin reorganization to transcriptional dysregulation in HGPS VSMCs. Our multi-omics resource provides unprecedented insights into how lamin A dysfunction epigenetically reprograms vascular cells, with implications for both HGPS and age-related cardiovascular disease.

While TADs remained globally stable across control and HGPS cells, we identified dynamic remodeling of chromatin states within TADs, especially involving H3K27me3 and H3K36me3 redistribution. Our observation that H3K27me3-enriched TADs were consistently the largest and least transcriptionally active is in line with the role of Polycomb-mediated repression in forming broad heterochromatic domains [29]. Notably, we detected a progressive shift from active (H3K36me3 or H3K27ac) to repressive (H3K27me3) chromatin states in a subset of TADs over replicative aging. This altered H3K27me3 landscape resembles the age-associated remodeling in facultative heterochromatin observed in multiple tissues [54–57], supporting the supposition that HGPS mirrors features of natural aging but with accelerated kinetics. To fully assess whether any stochastic component contributes to these changes will require single cell epigenomic profiling.

A/B compartmental dynamics describe the segregation of the genome into transcriptionally active (A) and inactive (B) domains, first identified through Hi-C mapping [58]. These compartments reflect underlying chromatin states, with A compartments enriched for euchromatic marks and B compartments associated with heterochromatin. Compartment organization is not static, it changes during development, differentiation, and disease. For example, it was demonstrated that dynamic compartment switching during stem cell differentiation, where genes transitioning from B to A become transcriptionally active [59]. In aging and age-related diseases, such as HGPS, these dynamics are disrupted, leading to compartment erosion and widespread gene misregulation [34,60]. Thus, A/B compartmental dynamics are essential for maintaining genome function and integrity, with their perturbation contributing to developmental defects and pathologies. We find that A/B compartmental dynamics further illustrate epigenome erosion in HGPS VSMCs. Early-passage HGPS VSMCs displayed compartment shifts with minimal transcriptional impact, whereas late passage cells showed stronger compartmental transitions associated with gene expression changes. Regions switching from A to B compartments showed increased H3K27me3 at late passages, reinforcing the concept that late-stage heterochromatin expansion contributes to transcriptional repression, as reported previously in HGPS fibroblasts and aging models [34,54,61]. Surprisingly, even regions strengthening their B compartment identity exhibited transcriptional activity, suggesting that structural compartment status alone is insufficient to enforce silencing, especially when repressive histone modifications are disrupted. This dissociation is consistent with reports showing that lamin-associated domains can become transcriptionally active during senescence or after lamina disassembly [56,61].

In addition to compartmental disruption, we observed a pronounced reduction in interchromosomal intermingling in HGPS VSMCs. Genome-wide analysis revealed that the majority of intermingling regions were specific to control cells, whereas HGPS cells displayed a substantial depletion of such contacts. Notably, both genes upregulated and downregulated in HGPS were preferentially associated with control-specific intermingling, indicating that transcriptional activation in HGPS frequently occurs despite the loss of these higher-order interactions. The stronger depletion of interchromosomal contacts surrounding downregulated genes further suggests that interchromosomal proximity may contribute to maintaining transcriptional competence in normal VSMCs. Together, these findings support a model in which premature aging is accompanied by erosion of interchromosomal genome organization, potentially destabilizing gene regulatory environments and contributing to widespread transcriptional dysregulation.

Importantly, we identified a global depletion of H3K27me3 within B compartments in HGPS cells. This depletion, despite preservation of structural compartmentalization, suggests a decoupling of spatial genome organization from epigenetic repression, and aligns with prior observations of PRC2 component downregulation and chromatin derepression in HGPS fibroblasts [34,56]. The redistribution of H3K27me3 from intergenic and intronic sites to promoter regions further underscores epigenetic drift as a key component of transcriptional instability in HGPS.

A significant body of research has established that the higher-order chromatin organization of the genome, from large-scale A/B compartments to sub-megabase TADs, is fundamental for regulating gene expression and maintaining cellular identity [21,58]. Disruptions to these broad architectural features have been widely implicated in cellular aging, senescence, and disease pathogenesis, including in premature aging syndromes like HGPS [34,62,63]. While these large-scale changes provide a crucial context for understanding epigenome-wide misregulation, they often fail to capture the specific molecular events that drive aberrant gene expression. The precise regulatory effects are often mediated by dynamic, high-resolution chromatin loops that bring distal enhancers into direct contact with their target promoters. A detailed analysis of these finer-scale interactions is therefore essential to fully elucidate the functional consequences of broader architectural disruptions. Beyond broad architectural disruptions, our loop-level analysis revealed that HGPS VSMCs accumulate disease-specific enhancer–promoter (E–P) loops that are tightly linked to vascular pathology gene expression. We observed consistent gain of expression among genes associated with HGPS-specific E–P loops, particularly at early passages, implicating enhancer rewiring as an early driver of aberrant gene activation. This is consistent with work in aging hematopoietic stem cells and cardiomyocytes, where altered enhancer connectivity precedes transcriptional reprogramming [64,65].

Disease-specific loops in HGPS were significantly enriched for genes involved in angiogenesis, extracellular matrix organization, and inflammatory signaling, processes central to atherosclerosis and vascular stiffening [38,40]. These findings suggest that chromatin rewiring in HGPS does not merely reflect epigenetic drift but actively promotes pathogenic gene networks. One striking example is GJA1, which gained a disease-specific loop and underwent transcriptional activation in HGPS, consistent with its role in promoting VSMC proliferation and intercellular signaling during atherosclerosis [50,51]. Conversely, at the PPARG locus, we observed the formation of a promoter–enhancer loop without corresponding transcriptional activation, due to persistent repressive chromatin features. This highlights that architectural contact alone is not sufficient for transcription, and must be interpreted in the context of local chromatin state and histone modification balance [66]. Furthermore, we observed increased CTCF and cohesin occupancy at loop anchors in HGPS, suggesting that aberrant recruitment of architectural proteins may contribute to non-canonical chromatin loops, in line with prior findings in cancer and aging models [63,67]. Whether this reflects compensatory stabilization of chromatin architecture or misregulated loop formation remains to be explored.

The redistribution of H3K27me3, particularly its accumulation at promoter regions and depletion from intergenic regions, reflects mislocalization of PRC2 machinery. Previous studies have linked this phenomenon to EZH2 misregulation in HGPS [34], and our findings reinforce this model by showing passage-dependent increases in promoter-proximal H3K27me3 despite transcriptional persistence at many loci. This local gain was in contrast with a global reduction of H3K27me3 in HGPS, suggesting PRC2 retargeting rather than complete loss. Consistent with a reduced density of H3K27me3 throughout the genome, polycomb group bodies were prominent in control VSMCs but disappeared in HGPS VSMCs. These are believed to be liquid condensates that form on facultative heterochromatin and are associated with the transcriptional repression of the genes contained within them. Since condensate formation is highly sensitive to local concentration, their absence is consistent with insufficient H3K27me3 density [68–70]. Support for such mutation-driven retargeting comes from cardiac laminopathy models, where the K219T mutation was found to increase the affinity of Lamin A/C for the PRC2 complex, this complex binds excessively to the *SCN5A* promoter, deposits repressive H3K27me3 marks, and sequesters the gene at the nuclear periphery, leading to transcriptional silencing [71]. Indeed, approximately 12% of promoters marked by new H3K27me3 sites in HGPS also retained H3K36me3, further indicating incomplete silencing [72,73]. The persistence of such regions may render these genes susceptible to erratic transcription, contributing to transcriptional noise and phenotypic heterogeneity, hallmarks of both HGPS and normative aging [31,74].

In summary, we report a progressive and multiscale collapse of chromatin regulation in HGPS VSMCs, characterized by widespread heterochromatin redistribution, compartmental drift and rewiring of enhancer–promoter loops. The dissociation between genome architecture and transcriptional control, evidenced by PPARG enhancer-promoter contact without activation, or B compartment strengthening without repression, suggests that chromatin misregulation in HGPS occurs not through singular loss of structure or function, but through complex epigenome disintegration. These findings have broad relevance beyond HGPS. Given the parallels between HGPS pathology and age-associated vascular disease, our study offers a framework for understanding how chromatin disorganization can drive VSMC dysfunction and atherosclerosis in aging populations. Therapeutically, interventions aimed at restoring the epigenetic landscape or enhancing PRC2 function may prove beneficial not only in HGPS but in aging-related vascular decline more generally [75,76].

This study focuses on VSMCs, limiting generalizability to other cell types in HGPS vasculopathy. Additionally, the *in vitro* model, while recapitulating progressive progerin accumulation, may not fully capture hemodynamic or immune contributions to atherosclerosis. Future work should explore cell-type-specific chromatin dynamics in aged or vascular disease tissue or from tissue-engineered HGPS arteries and test interventions targeting epigenetic regulators such as EZH2 to restore chromatin homeostasis.

## 4 Methods

### Resources and cell culture

iPSC lines were previously described [17]. Cell lines 0901C and 0031C are available from the Progeria Research Foundation under an MTA; BJ1C and AG1B are available from the corresponding author (WLS). RNA-seq data are deposited in GEO (GSE231761) [18]. All iPSC lines were cultured on Matrigel-coated plates in E8 medium (DMEM/F12 supplemented with L-ascorbic acid, sodium selenite, βFGF, TGF-β1, sodium bicarbonate, holo-transferrin, and gentamicin) as previously described [17].

### VSMC differentiation

iPSCs were differentiated into VSMCs following a previously described protocol [77] with minor modifications. Briefly, embryoid bodies (EBs) were generated by treating iPSCs with Collagenase IV (Life Technologies) for 30 min at 37 °C and gently dislodging them with a cell scraper. Cell clusters were transferred to low-attachment plates (Corning) and cultured in EB medium (DMEM/F12 with 20% Knockout Serum Replacement [Invitrogen], GlutaMax, β-mercaptoethanol, nonessential amino acids, and gentamicin) for 7–10 days. EBs were then plated on 0.1% gelatin-coated plates for 3–5 days, after which outgrowths were trypsinized and replated on Matrigel in Medium 231 with growth supplement. Cells were passaged at ∼80% confluency. For VSMC differentiation, cells were plated on gelatin-coated 6-well plates and cultured in Medium 231 with differentiation supplement (Life Technologies) for 7 days, with characterization performed at each passage.

### In situ Hi-C

In situ Hi-C was performed as previously described, with modifications [78]. Briefly, two cell lines of control and HGPS VSMCs (8 × 10^6 cells) were crosslinked in 1% formaldehyde for 10 min at room temperature and quenched with 0.2 M glycine. Crosslinked cells were pelleted by centrifugation and stored at −80 °C. Nuclei were isolated and chromatin was digested overnight with 100 U of DpnII (NEB). Following digestion, 5′ overhangs were filled in with biotin-14-dATP (Life Technologies) and blunt-end ligation was performed in situ for 4h. Crosslinks were reversed by proteinase K treatment, and DNA was purified by phenol:chloroform extraction, ethanol precipitation, and resuspension in low EDTA, tris-buffered water (TLE). Library quality was assessed as described previously [78]. A known chromatin interaction was amplified by PCR, and the product was subjected to differential digestion with ClaI and MboI to confirm efficient biotin incorporation. For library preparation, 40 µg of DNA from each sample was fragmented to 200–300 bp using a sonicator (Covaris). Fragmented DNA was size-selected with AMPure XP beads (Beckman Coulter) according to the manufacturer’s instructions. The purified DNA fragments underwent end repair, after which biotinylated DNA was isolated using Streptavidin beads. Subsequently, polyA tailing and adapter ligation (Illumina TruSeq) were performed to generate the final libraries, and libraries were PCR-amplified and purified. Final libraries were sequenced on the NovaSeq 6000 platform (Illumina) with 150 bp paired-end reads to produce over 500M read pairs at the Donnelly Sequencing Centre (Toronto). Hi-C data was processed using the tools and workflow from HiCExplorer [79]. Fastq files were aligned to Homo sapiens GRCh38 primary assembly (from GENCODE) using BWA-MEM version 0.7.17-r1188 (https://github.com/lh3/bwa) with the following parameters: -A1 -B4 -E50 -L0 -t 20. Hi-C matrices for replicates were built using hicBuildMatrix at 5kb resolution and merged with hicSumMatrix to produce combined matrices for control and HGPS VSMCs samples at passages 7 and 14. Conversions between Hi-C formats, depending on the downstream analysis, were performed using hicConvertFormat tool.

### CUT&Tag

Cleavage Under Targets and Tagmentation (CUT&Tag) was performed essentially as described [80], with minor modifications using biological triplicates. Briefly, three cell lines of control and HGPS VSMC cells (100,000) were harvested, washed twice in Wash Buffer (20 mM HEPES pH 7.5, 150 mM NaCl, 0.83 mM spermidine, 0.1% BSA, Roche Protease inhibitor cocktail). The CUT&Tag-IT® Spike-In Control (Active Motif), composed of Drosophila nuclei for downstream normalization and quantitative comparison, was added to each sample at a 1:20 ratio of spike-in to cells, and bound to concanavalin A–coated magnetic beads (Bangs Laboratories). Bead-bound cells were permeabilized in Dig-Wash Buffer (Wash Buffer supplemented with 0.05% digitonin) and incubated with primary antibody (1:100 dilution, overnight at 4 °C with rotation). After three washes, samples were incubated in an appropriate secondary antibody (1:100 dilution, 1 h at room temperature). Beads were washed and incubated with pA–Tn5 transposome (Epicypher, pre-loaded with sequencing adapters) for 1 h at room temperature. After additional washes in Dig-300 Buffer (0.05% Digitonin, 20 mM HEPES, pH 7.5, 300 mM NaCl, 0.83 mM Spermidine, Roche Protease inhibitor cocktail), targeted tagmentation was initiated by the addition of MgCl₂-containing tagmentation buffer (10 mM MgCl₂ in Dig-300 Buffer) for 1 h at 37 °C. DNA was released by incubation in 0.1% SDS, 16 mM EDTA at 50 °C for 1 h with proteinase K digestion. DNA was purified using phenol:chloroform and libraries were amplified by PCR (12–15 cycles) with Q5 Hot Start High-Fidelity DNA Polymerase (New England Biolabs) using dual-indexed barcoded primers. Amplified libraries were sent for further processing and sequencing at IRCM, Montreal. Libraries were sequenced on an Illumina NovaSeq 6000 platform with paired-end 100 bp reads. Sequencing data were processed using the nf-core cutandrun pipeline version 3.2.2 (doi: 10.5281/zenodo.5653535). Peak calling for all CUT&Tag data was performed using SEACR (Sparse Enrichment Analysis for CUT&RUN) version 1.3 with parameters: target.bedgraph IgG.bedgraph norm stringent output. Peak replicates were merged into a consensus set of peaks per sample using bedtools version 2.31.0 [81] Genomic distribution and annotation of H3K27me3, H3K36me3 and H3K27ac peaks were performed using ChIPseeker version 1.38.0 [82]. All CUT&Tag signal tracks in bigWig format were generated using deeptools version 3.5.0 [83] bamCoverage with RPGC normalization. Signal enrichment analysis of CTCF at TAD and loop boundaries was computed using deeptools computeMatrix and plotHeatmap. Likewise, in the case of histone mark H3K27me3 signal across gene bodies.

We used the following antibodies: Histone H3K27me3 (NEB, 9733T), Guinea Pig Anti-rabbit IgG, Rabbit Anti-mouse IgG (Antibodies-online, ABIN101961), Histone H3K36me3 (Active Motif, 61022), Histone H3K27ac (ab177178), CTCF (EMD-Millipore, 07-729-25UL), SMC1A (ab9262), Spike-In Control Anti-rabbit (Active Motif, 53168), and Spike-In Control Anti-mouse (Active Motif, 53173).

### DNA-FISH

#### Cells and fixation

HGPS VSMCs were grown on No. 1.5 glass coverslips to 70–80% confluency. Cells were fixed in 4% paraformaldehyde in PBS for 10 min at room temperature, washed three times in PBS, and permeabilized in 0.2% Triton X-100 in PBS for 10 min at room temperature. Coverslips were transferred to 70% ethanol and stored overnight at 4 °C.

#### Pre-treatments for 3D preservation and access

Coverslips were rehydrated through 50% ethanol to PBS, then incubated in 0.1 N HCl for 10 min at room temperature, rinsed in PBS, and equilibrated in 2x SSC. RNase A treatment was performed separately (100 µg/mL RNase A in 2x SSC, 37 °C for 45 min), followed by 2x SSC washes and equilibration in 50% formamide/2x SSC (pH ∼7.0) for ≥30 min at room temperature.

#### Probe design and pools

Single-stranded DNA oligos were designed with OligoMiner principles and ordered at 25 nmol scale, standard desalting. Each targeting oligo comprised a 5′ universal priming handle (20 nt), a TTTT spacer, and a 30–37 nt genomic homology arm (total length ∼50–57 nt). Reporter oligos complementary to the 5′ handles were synthesized with 5′ dyes: FAM for Region 1, Cy3 for Region 2, and Cy5 for Region 3. For this study, we used three color-coded probe sets per locus: PPARG: three pools (Region 1/2/3), 10 oligos per region (30 total), 50–57 nt each. GJA1: three pools (Region 1/2/3), 10 / 10 / 8 oligos, 50–57 nt each. Oligo names and sequences are contained in the IDT order sheet (Table S5); pools were mixed by region for single-step hybridization.

#### Hybridization mix and co-denaturation

Hybridization was performed in a 50 µL volume per 18 x 18 mm coverslip using 50% formamide, 2x SSC, 10% dextran sulfate, 1% Tween-20, with salmon sperm DNA (100 µg/mL) as competitor. Probe pools were combined to a final pool concentration of 100 nM per color (total concentration across tiles within a pool). Probes were heat-denatured at 70 °C for 5 min and snap-cooled on ice immediately before use. For 3D-preserving co-denaturation, coverslips were mounted with 50 µL hybridization mix, sealed, and heated to 75 °C for 3 min, then snap-cooled and incubated overnight at 37 °C in a humidified chamber.

#### Post-hybridization washes and mounting

Coverslips were washed four times for 3 min in 2x SSC at 45 °C, then four times for 3 min in 0.1x SSC at 60 °C. After a brief rinse in 2x SSC at room temperature, samples were stained with DNA dye Hoechst and mounted with Mowiol before proceeding to imaging.

#### Imaging

3D stacks were acquired on a Zeiss AxioImager upright epifluorescence microscope with a 100x/1.4 NA oil objective. Approximately 50 z-slices were collected at 0.20 µm axial step size with identical exposure settings and channel order for all conditions. Images were saved as 16-bit stacks with calibrated voxel sizes; exported datasets had effective voxel sampling of around 0.065 x 0.065 x 0.30 µm.

#### Image analysis

3D analysis was performed in Imaris. Nuclei (Hoechst) were segmented into surfaces using intensity thresholds with size filtering to exclude debris/blebs. FISH foci were segmented per channel as separate surfaces with local background subtraction and a minimum-size filter to remove noise. For each cell, we calculated: (i) centroid-to-centroid distances among Region 1/2/3 within the locus, (ii) distances from each focus to the nuclear centroid and boundary, and (iii) inter-allelic distances where applicable. At least 100 cells per biological replicate were analyzed across 3 replicates per condition (n∼ 300 cells/condition). Cells with obvious flattening, segmentation artifacts, or out-of-focus stacks were excluded a priori.

### Immunofluorescence staining

Human HGPS VSMCs were seeded on Matrigel-coated Greiner µClear 96 well plates and cultured to 80% confluency. Cells were fixed in 4% paraformaldehyde in PBS for 10 min at room temperature, then washed three times in 1x PBS. Permeabilization was performed with 0.5% Triton X-100 in PBS for 10 min at room temperature, followed by three PBS washes. Non-specific binding was blocked with 3% BSA in PBS for 30 min at room temperature and washed three times in PBS. Primary antibodies were applied in the blocking buffer for 45 min at room temperature: Rabbit anti-EZH2 (Active Motif 39639 1:200). After incubation, wells were rinsed once with 0.1% Triton X-100 in PBS, then washed three times in PBS (1 min each). Fluorophore-conjugated secondary antibody was added in a blocking buffer for 45 min at room temperature: goat anti-rabbit Cy3 (Jackson ImmunoResearch 111-165-144, 1:300), followed by three PBS washes. DNA was counterstained with Hoechst 33342 for 20 min at room temperature. Plates were covered with Mowiol 4-88 mounting medium (Calbiochem 475904) prior to imaging. Images were acquired on a high-content imaging system (ImageXpress XLS, Molecular Devices) controlled by MetaXpress software using an sCMOS camera and a 20x/0.75 NA Plan Apo objective. Exposure and illumination settings were held constant across conditions. For each condition, multiple fields were collected to yield around 1,000–4,000 cells. Quantification was performed in CellProfiler 4.2.8. After per-image background subtraction, nuclei were segmented from the Hoechst channel to generate nuclear masks. For each nucleus, integrated intensity (sum of pixel values) of EZH2 channels was measured. Data from all fields per condition were pooled for downstream analysis.

### Identification and classification of TADs

Comparison of Hi-C matrices counts enrichment was computed using HiCExplorer tools. Matrices binned at 50kb were first normalized using Imakaev’s iterative correction (ICE) via hicCorrectMatrix, and the distance vs count relationship was visualized with hicPlotDistVsCounts. TAD calling was performed using SpectralTAD version 1.18.0 [84] at 100kb resolution with Vanilla-Coverage (VC) as normalization method. Analysis of TAD chromatin states was performed by creating a matrix of H3K27ac, H3K36me3, and H3K27me3 signal across TADs using deeptools multiBigwigSummary BED-file. TAD classification was assigned based on the histone mark with the highest peak signal, contingent on that signal being within the top 25% of its overall distribution. TADs that did not meet these conditions were classified as “other TAD”. Differences between chromatin state annotated TADs were analyzed, such as TAD sizes, gene density using bedtools intersect, and gene expression at these annotated TADs using TPM counts from RNA-Seq (GSE231761) [18]. Statistical differences between TAD states were determined using the Mann-Whitney-Wilcoxon test, with an adjusted p-value < 0.05 considered to be statistically significant. Transitions in TAD chromatin states were analyzed for common TADs (with identical coordinates) between conditions, both between samples (Control vs. HGPS VSMCs) and across passages (Control P7 vs. P14). These transitions were visualized using alluvial plots with the R package ggalluvial.

### Characterization of loops

Loop calling and differential loop detection were computed with Mustache version 1.2.0 [85] at 5kb resolution. Chromatin loops were linked to promoters (upstream 1kb of genes) and GeneHancer [86] enhancers using R package LoopRig version 0.1.1 (10.32614/CRAN.package.LoopRig) LinkedElements function. Analysis of gene expression changes in loops and differential loops anchored to E-P (enhancers and promoters) was computed by filtering for those genes that were differentially expressed (log2 fold change cutoff > 0 and adjusted p-value threshold at 0.05; HGPS vs control) between samples from RNA-seq data. Statistical significance was calculated using Mann-Whitney-Wilcoxon test, with a p-value < 0.05 considered statistically significant. Gene ontology and KEGG pathway [87] enrichment analysis of genes that fall under differential loops anchored by E-P were computed using g:Profiler web server [88]. Visualization of differential loops on Hi-C contact map (5kb resolution) at genes of interest was performed using Juicebox [89] by loading HGPS and control matrices using the “Observed vs Control” feature, splitting both maps across the diagonal.

### Analysis of A/B compartment transitions

A/B compartments and differential compartments were identified using dcHiC version 2.1 [33] at 50kb resolution with default parameters. To investigate changes in gene expression between compartment transitions, TPM counts from RNA-Seq were used by overlapping genes at differential compartments (binned at 50kb). Statistical significance was calculated using Wilcoxon Signed-Rank test, with p-value < 0.05 as statistically significant. Differential compartments were categorized as transitions (A-to-B or B-to-A) if their compartment scores changed sign, or as matches (A-to-A or B-to-B) if the scores remained the same. Those compartments that didn’t change sign (A-to-A or B-to-B) were further classified into strength changes based on compartment scores being higher in control or in HGPS VSMCs. The transitions were visualized using alluvial plots. Additionally, A/B compartment classification was further validated by assessing their correlation with active and repressive histone marks H3K27ac and H3K27me3, respectively, through manual inspection of signal tracks using Integrative Genomics Viewer (IGV) [90]. To assess changes in H3K27me3 at compartment switch regions, we calculated the log2 ratio of H3K27me3 signal between HGPS and control VSMCs using deeptools bigwigCompare and then quantified these ratios at A-to-B and B-to-A switches with multiBigwigSummary. Similarly, differences in global enrichment of H3K27me3 signal at all B compartments between HGPS and control were also examined using deeptools multiBigwigSummary. For both analyses, statistical significance was calculated using Mann-Whitney-Wilcoxon test, with a p-value < 0.05 considered statistically significant.

### Interchromosomal intermingling analysis

Hi-C intermingle analysis was performed by generating interchromosomal matrices at 250kb resolution using Juicer tools [91]. Comparative analyses of the matrices, including visualization and quantitative assessment, were performed in Python version 3.12.8 following the procedure as described in [92].

### Visualization

For integrated visualization of A/B compartments, TADs, loops and bigWig tracks of histone marks/proteins, pyGenomeTracks version 3.9 [93] was used to produce genome browser tracks of all the datasets at regions of interest.

## Data availability

All genomics data are available via GEO: CUT&Tag datasets (GSE312029), Hi-C datasets (GSE312031), and as well as previously published RNA-seq datasets (GSE231761).

## Author Contributions

**Conceptualization:** M.N., M.J.H., T.J.P, and W.L.S.; **Methodology:** M.N., R.K., and A. S.; **Analysis:** M.N., N.A., R.K.; **Writing:** M.N., N.A., R.K., M.J.H.; **Editing:** All authors contributed to editing the manuscript. **Funding:** M.J.H., T.J.P, and W.L.S.

## Acknowledgements

This work was supported by the Canadian Institutes of Health Research [PJT-178364] to WLS, TJP, and MJH. We would like to acknowledge the following support: Dr. Xuejun Sun and the Cell Imaging Facility at the Cross Cancer Institute for instrument and analysis assistance and Jordan Brooks for technical assistance, the Ottawa Human Pluripotent Stem Cell Facility (RRID:SCR_027437), the OHRI/uOttawa Bioinformatics Core (RRID:SCR_022466), the OHRI High Content Imaging Core, and StemCore (RRID:SCR_012601). MJH is supported by a Tier 1 Canada Research Chair in Genome Cell Biology and Dynamics. WLS has been supported by the University of Ottawa Distinguished Research Chair in Disease Modeling and Therapeutic Discovery and a Tier 1 Canada Research Chair in Integrative Stem Cell Biology.

## Ethics declarations

The induced pluripotent stem cell lines [17,18] were generated in collaboration with the Progeria Research Foundation under the approval of the Stem Cell Oversight Committee and the Ottawa Hospital Research Ethics Board.

## Supplemental Figure legends

**Fig. S1:**
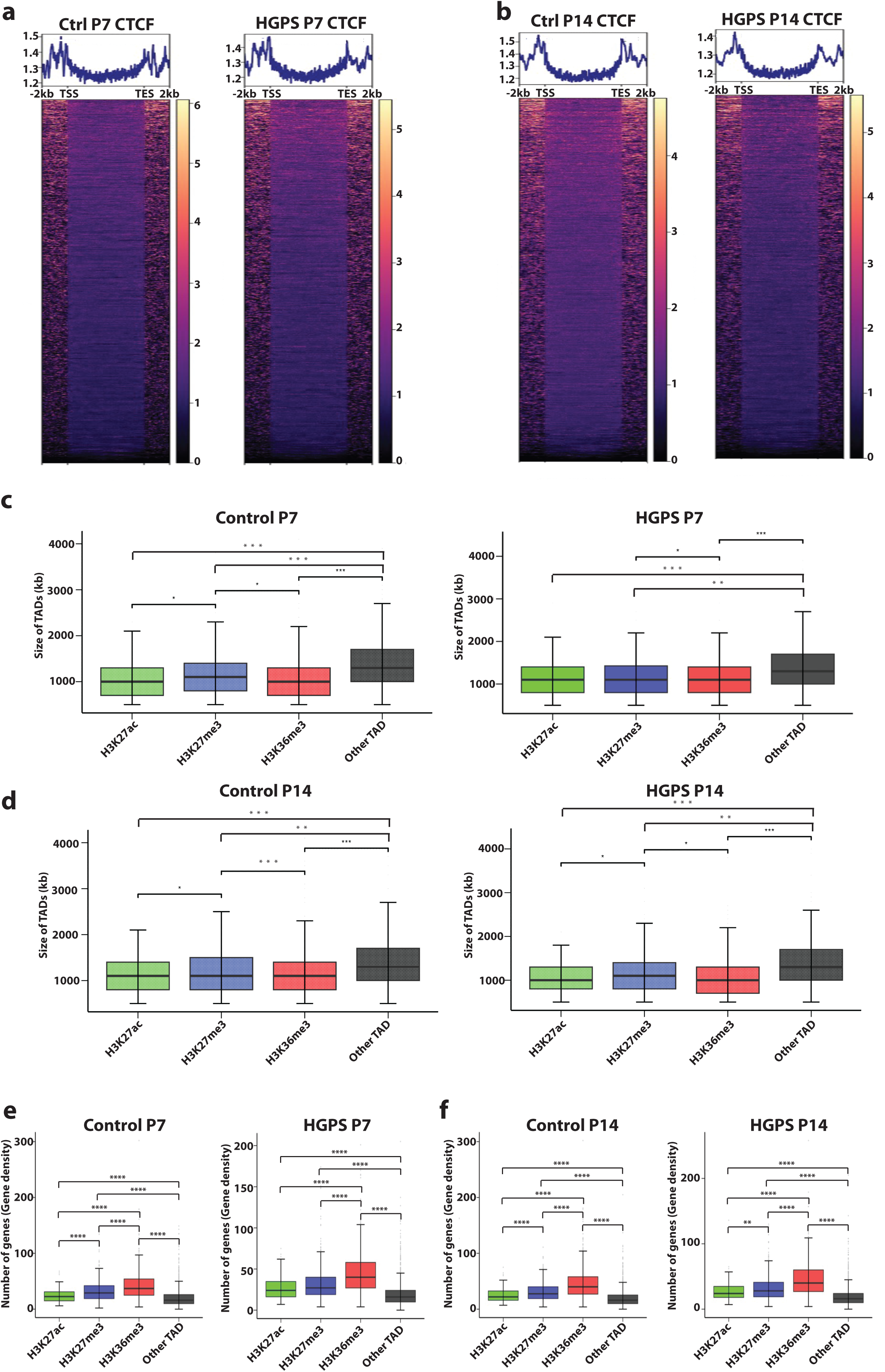
CTCF binding profiles at TAD boundaries reveal conserved boundary enrichment in control and HGPS VSMCs. **a** Heatmap of normalized CTCF coverage around TADs between control and HGPS VSMCs at P7 and P14. The signal intensity is measured in log2 read coverage normalized by sequencing depth. **b, c** Size of histone mark-defined chromatin states of TADs for control and HGPS samples. Panel **b** shows samples at passage 7, while panel **c** is at passage 14. **d** Gene density of TAD chromatin states between control and HGPS VSMCs at P7 and P14. P-values are computed using the Mann-Whitney-Wilcoxon test, *adj. p value < 0.05, **adj. p value < 0.01, ***adj. p value < 0.001.

**Fig. S2:**
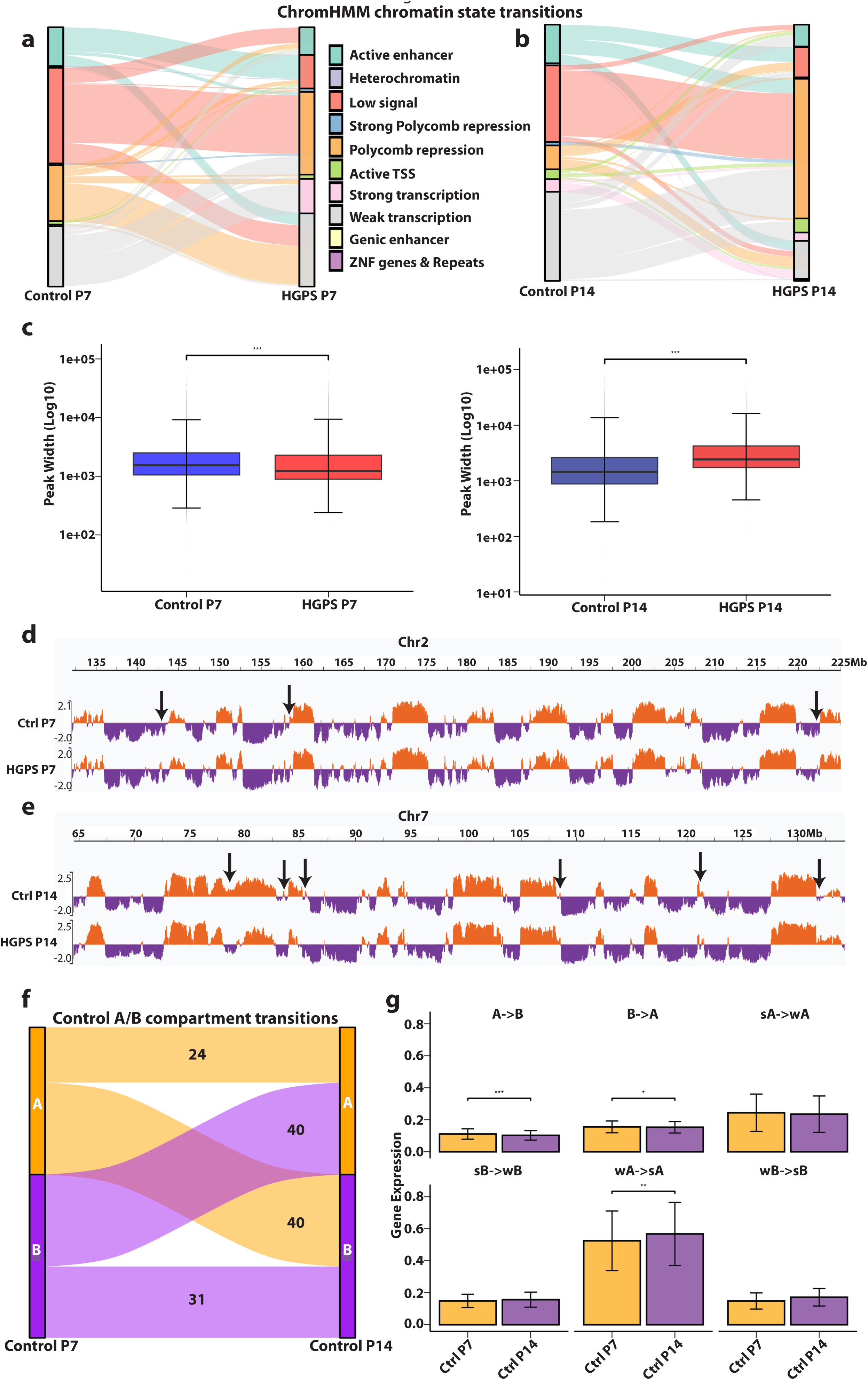
**a** ChromHMM chromatin state transitions between common TADs in Control and HGPS, annotated using pre-trained 18 state model. **b** Same as (**a**) but for P14**. c** Quantification of H3K27me3 peak width (log10) at H3K27me3-enriched TADs between control and HGPS VSMCs at P7 and P14, suggesting spreading of H3K27me3 domains. Representative IGV genome tracks at chr2:132,121,064-228,998,499 (**d**) and chr7:64,498,754-144,171,740 (**e**) regions showing differential A/B compartments between control and HGPS VSMCs. A compartments are represented in orange, while B compartments are in purple. Arrows show differential A/B compartments. **f** Differential A/B compartment transitions between control VSMCs P7 and control VSMCs P14 at 50kb resolution (A compartments are represented in orange, while B compartments are in purple). **g** Gene expression at compartment transitions between control VSMCs P7 and control VSMCs P14. Strong (s) and weak (w) compartments were classified by comparing the relative compartment strength between samples. P-values are computed using the Mann-Whitney-Wilcoxon test, *adj. p value < 0.05, **adj. p value < 0.01, ***adj. p value < 0.001.

**Fig. S3:**
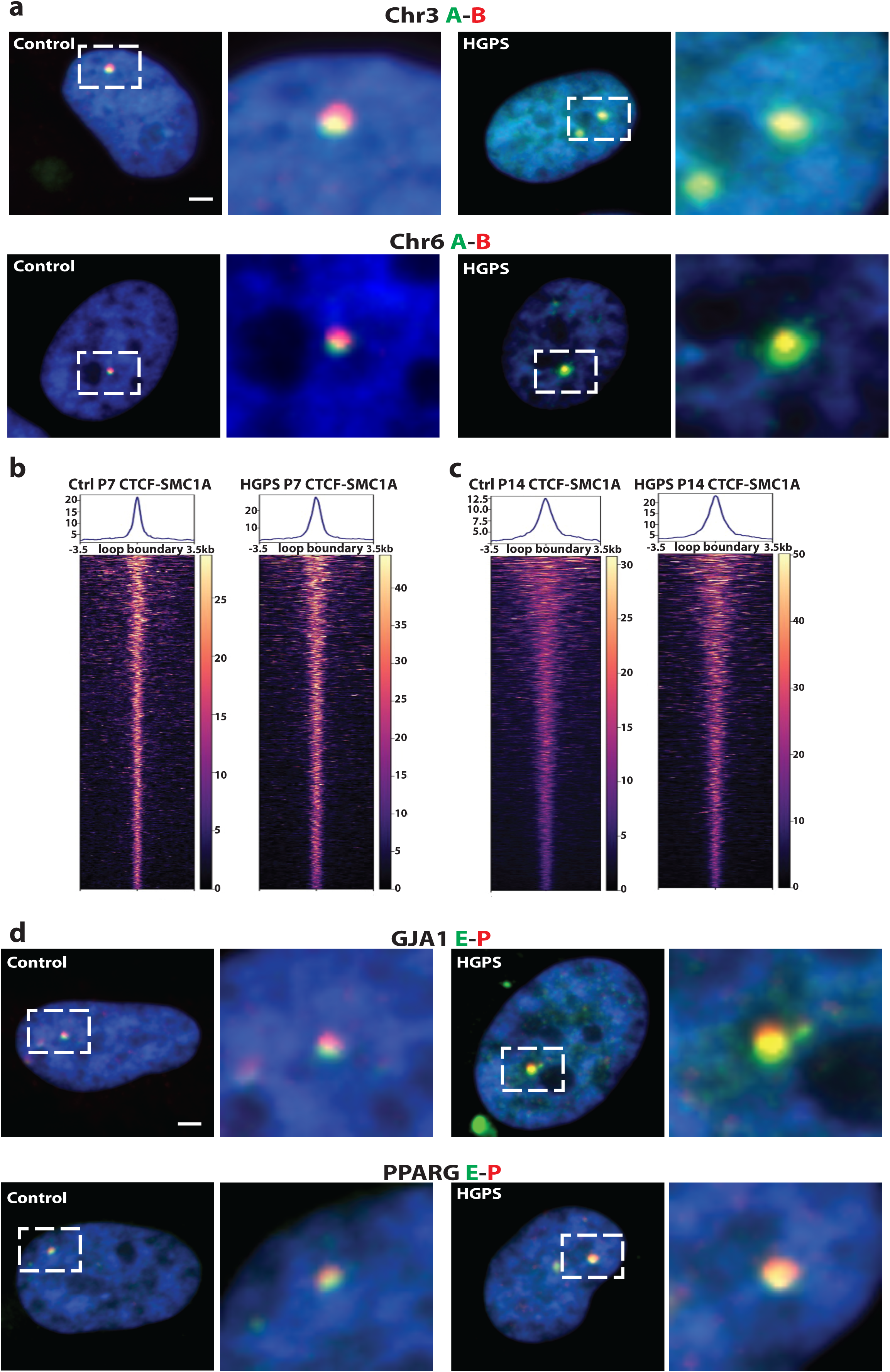
**a** Representative DNA-FISH images of chromosome 3 and chromosome 6 loci in control and HGPS VSMCs. Probes A (green) and B (red) are spatially distinct in control nuclei but exhibit significant overlap (yellow) in HGPS, indicating a localized collapse of 3D chromatin architecture. Insets show magnification of boxed regions. Scale bar: 2 µm. Heatmap of normalized CTCF coverage overlapping with SMC1A covering 3.5 kb up-and downstream of CTCF binding sites at the loop boundary between control and HGPS VSMCs at P7 (**b**) and P14 (**c**). The signal intensity is measured in log2 read coverage normalized by sequencing depth. **d** Representative DNA-FISH images of *GJA1* and *PPARG* loci. Probes target the enhancer (E, green) and promoter (P, red) regions. Increased signal overlap (yellow) in HGPS nuclei indicates spatial proximity between these regulatory elements compared to control cells. Insets show magnified views of boxed regions. Scale bar: 2 µm

**Fig. S4:**
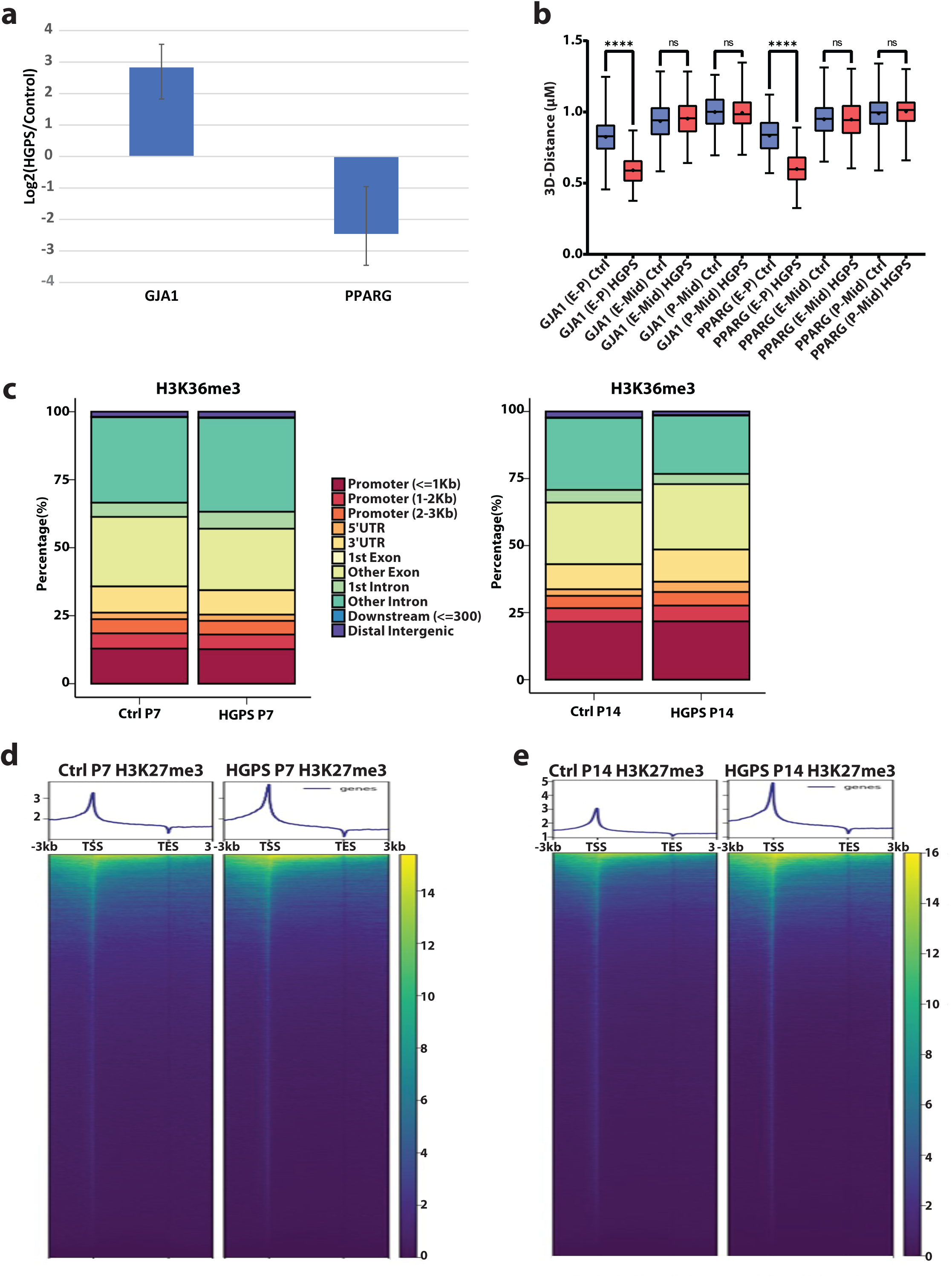
**a** PPARG and GJA1 transcript levels assessed by qPCR. **b** PPARG and GJA1 enhancer-promoter (E-P) loops validated by DNA-FISH. Enhancer-midpoint (E-Mid) and promoter-midpoint (P-Mid) probes were used to test for sensitivity of the signal**. c** Genomic distribution of H3K36me3 peaks for control and HGPS at P7 (left panel) and P14 (right panel). Heatmap representation of H3K27me3 distribution across gene bodies, 3kb up or downstream of transcription start site (TSS) and transcription end site (TES) at passage 7 (**d**) and passage 14 (**e**). H3K27me3 is redistributed to gene TSS in HGPS VSMCs late passage. P-values are computed using the Mann-Whitney-Wilcoxon test, *adj. p value < 0.05, **adj. p value < 0.01, ***adj. p value < 0.001.

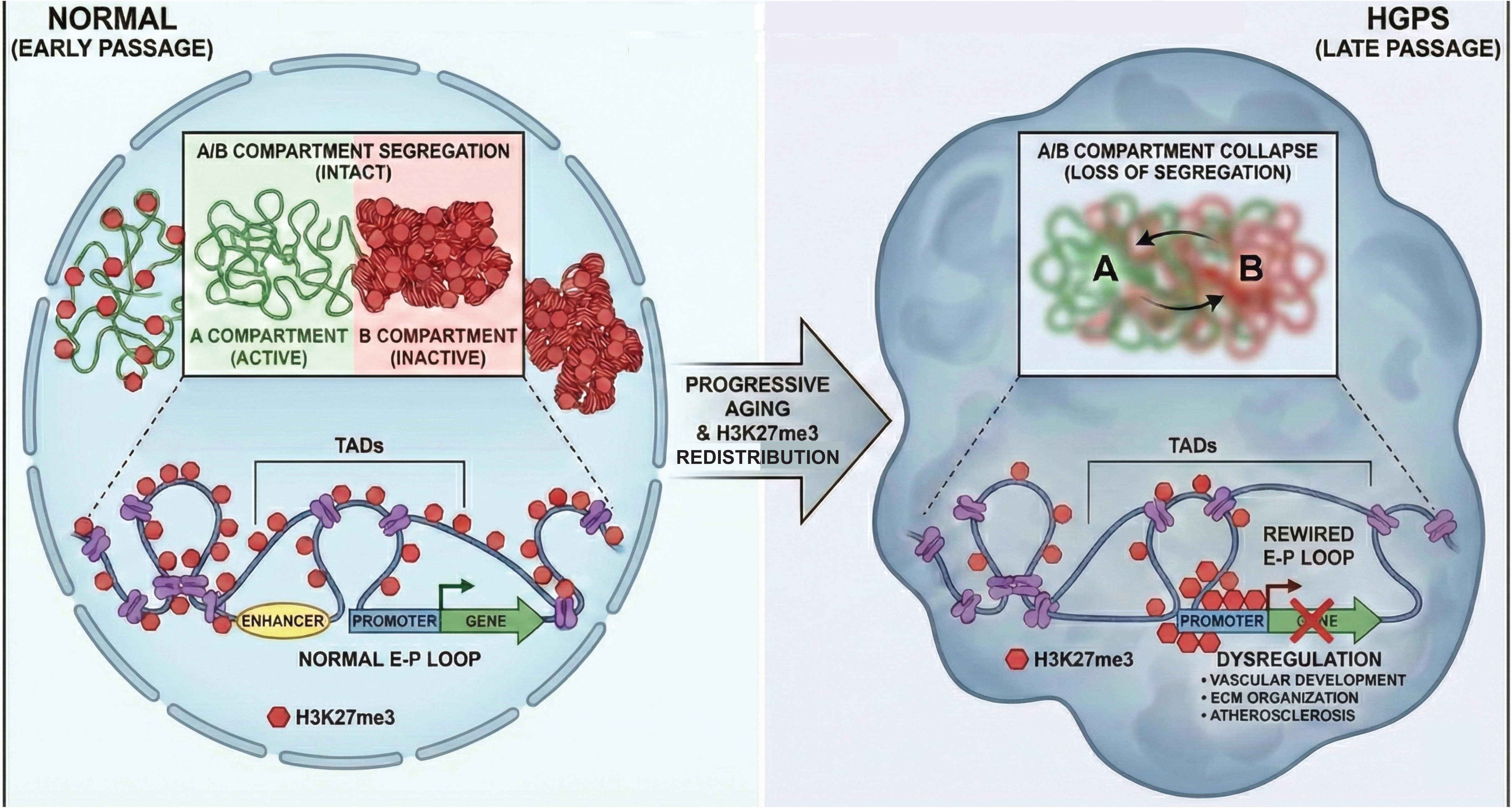

## Supplemental tables

**Table S1a:**
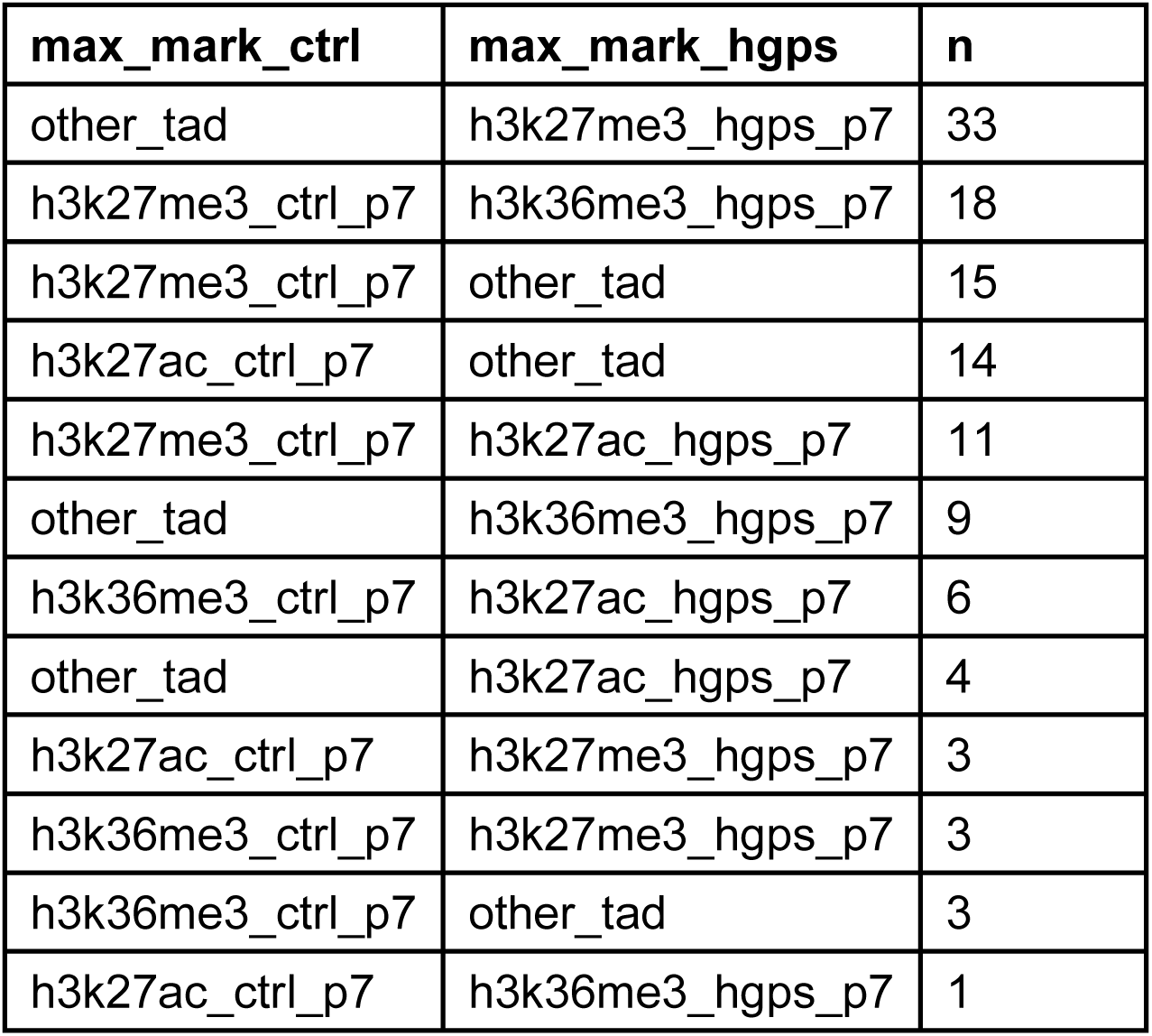
TAD chromatin state transition from Control P7 to HGPS P7.

**Table S1b:** TAD chromatin state distribution at Control P7.

| max_mark | n | fraction |
| --- | --- | --- |
| other_tad | 1340 | 0.58746 |
| h3k27ac_ctrl_P7 | 92 | 0.04033 |
| h3k36me3_ctrl_P7 | 417 | 0.18282 |
| h3k27me3_ctrl_P7 | 432 | 0.18939 |

**Table S1c:** TAD chromatin state distribution at HGPS P7.

| max_mark | n | fraction |
| --- | --- | --- |
| other_tad | 1322 | 0.58161 |
| h3k27ac_hgps_P7 | 102 | 0.04487 |
| h3k36me3_hgps_p7 | 425 | 0.18698 |
| h3k27me3_hgps_P7 | 424 | 0.18654 |

**Table S2a:** TAD chromatin state transition from Control P14 to HGPS P14.

| max_mark_ctrl | max_mark_hgps | n |
| --- | --- | --- |
| h3k36me3_ctrl_p14 | h3k27me3_hgps_p14 | 33 |
| h3k36me3_ctrl_p14 | h3k27ac_hgps_p14 | 32 |
| other_tad | h3k27me3_hgps_p14 | 30 |
| h3k27ac_ctrl_p14 | other_tad | 18 |
| h3k27me3_ctrl_p14 | other_tad | 18 |
| h3k27ac_ctrl_p14 | h3k27me3_hgps_p14 | 5 |
| h3k36me3_ctrl_p14 | other_tad | 4 |
| other_tad | h3k27ac_hgps_p14 | 4 |
| h3k27ac_ctrl_p14 | h3k36me3_hgps_p14 | 2 |
| h3k27me3_ctrl_p14 | h3k27ac_hgps_p14 | 2 |
| other_tad | h3k36me3_hgps_p14 | 2 |

**Table S2b:** TAD chromatin state distribution at Control P14.

| max_mark | n | fraction |
| --- | --- | --- |
| other_tad | 1268 | 0.56785 |
| h3k27ac_ctrl_P14 | 127 | 0.05687 |
| h3k36me3_ctrl_p14 | 444 | 0.19884 |
| h3k27me3_ctrl_P14 | 394 | 0.17644 |

**Table S2c:** TAD chromatin state distribution at HGPS P14.

| max_mark | n | fraction |
| --- | --- | --- |
| other_tad | 1298 | 0.56855 |
| h3k27ac_hgps_P14 | 150 | 0.0657 |
| h3k36me3_hgps_p14 | 354 | 0.15506 |
| h3k27me3_hgps_P14 | 481 | 0.21069 |

**Table S3:** TAD chromatin state transition from Control P7 to Control P14.

| max_mark_ctrl_P7 | max_mark_ctrl_P14 | n |
| --- | --- | --- |
| other_tad | h3k27me3_ctrl_p14 | 46 |
| h3k27me3_ctrl_p7 | h3k36me3_ctrl_p14 | 27 |
| h3k27me3_ctrl_p7 | other_tad | 23 |
| other_tad | h3k27ac_ctrl_p14 | 23 |
| other_tad | h3k36me3_ctrl_p14 | 11 |
| h3k27me3_ctrl_p7 | h3k27ac_ctrl_p14 | 9 |
| h3k27ac_ctrl_p7 | other_tad | 8 |
| h3k36me3_ctrl_p7 | h3k27ac_ctrl_p14 | 6 |
| h3k36me3_ctrl_p7 | other_tad | 4 |
| h3k27ac_ctrl_p7 | h3k36me3_ctrl_p14 | 3 |

**Table S4:** TAD chromatin state transition from HGPS P7 to HGPS P14.

| max_mark_hgps_P7 | max_mark_hgps_P14 | n |
| --- | --- | --- |
| h3k36me3_hgps_p7 | h3k27me3_hgps_p14 | 33 |
| h3k36me3_hgps_p7 | h3k27ac_hgps_p14 | 27 |
| h3k27me3_hgps_p7 | other_tad | 23 |
| other_tad | h3k27me3_hgps_p14 | 21 |
| other_tad | h3k27ac_hgps_p14 | 13 |
| h3k27ac_hgps_p7 | other_tad | 6 |
| h3k36me3_hgps_p7 | other_tad | 4 |
| h3k27ac_hgps_p7 | h3k36me3_hgps_p14 | 3 |
| h3k27ac_hgps_p7 | h3k27me3_hgps_p14 | 2 |
| h3k27me3_hgps_p7 | h3k36me3_hgps_p14 | 2 |
| other_tad | h3k36me3_hgps_p14 | 2 |

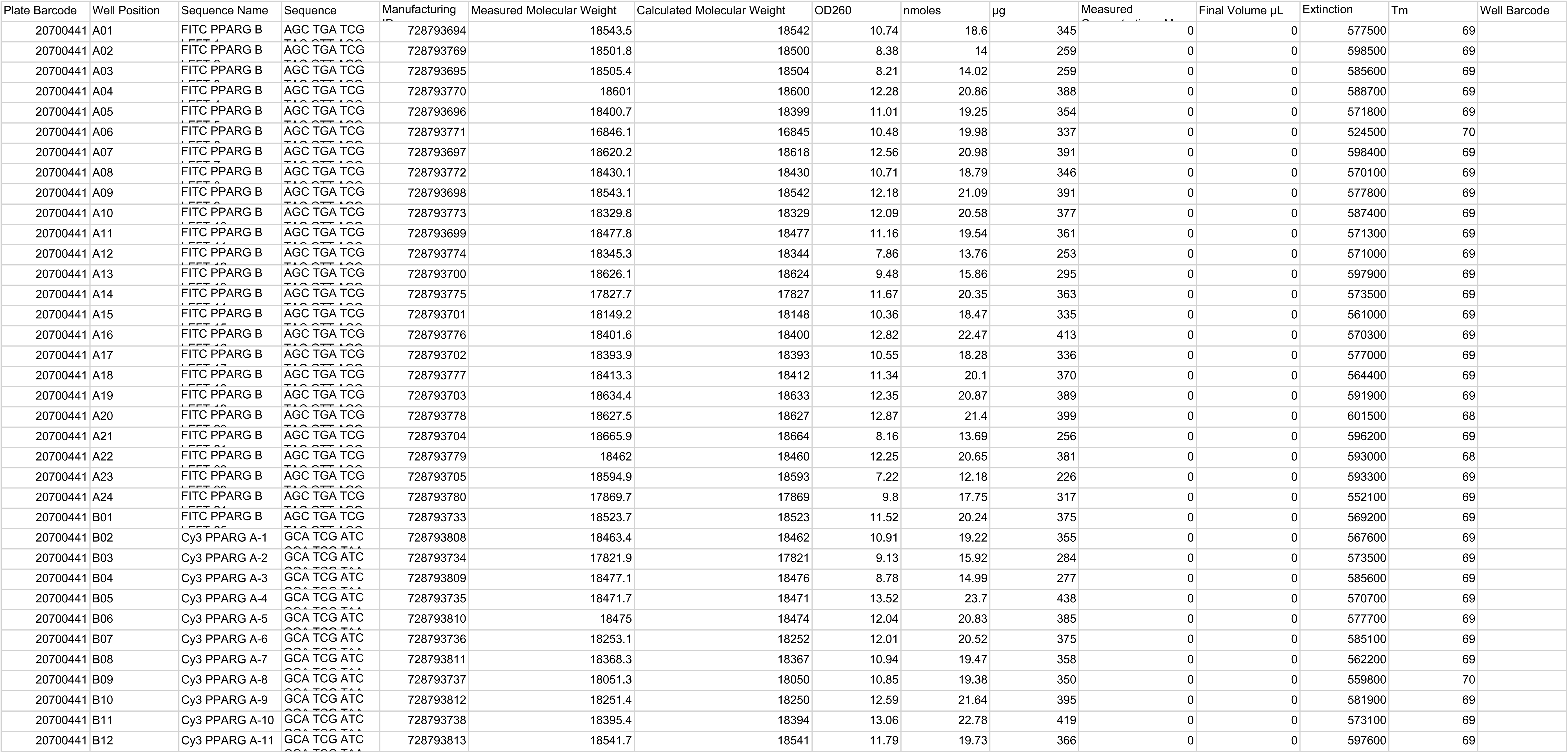
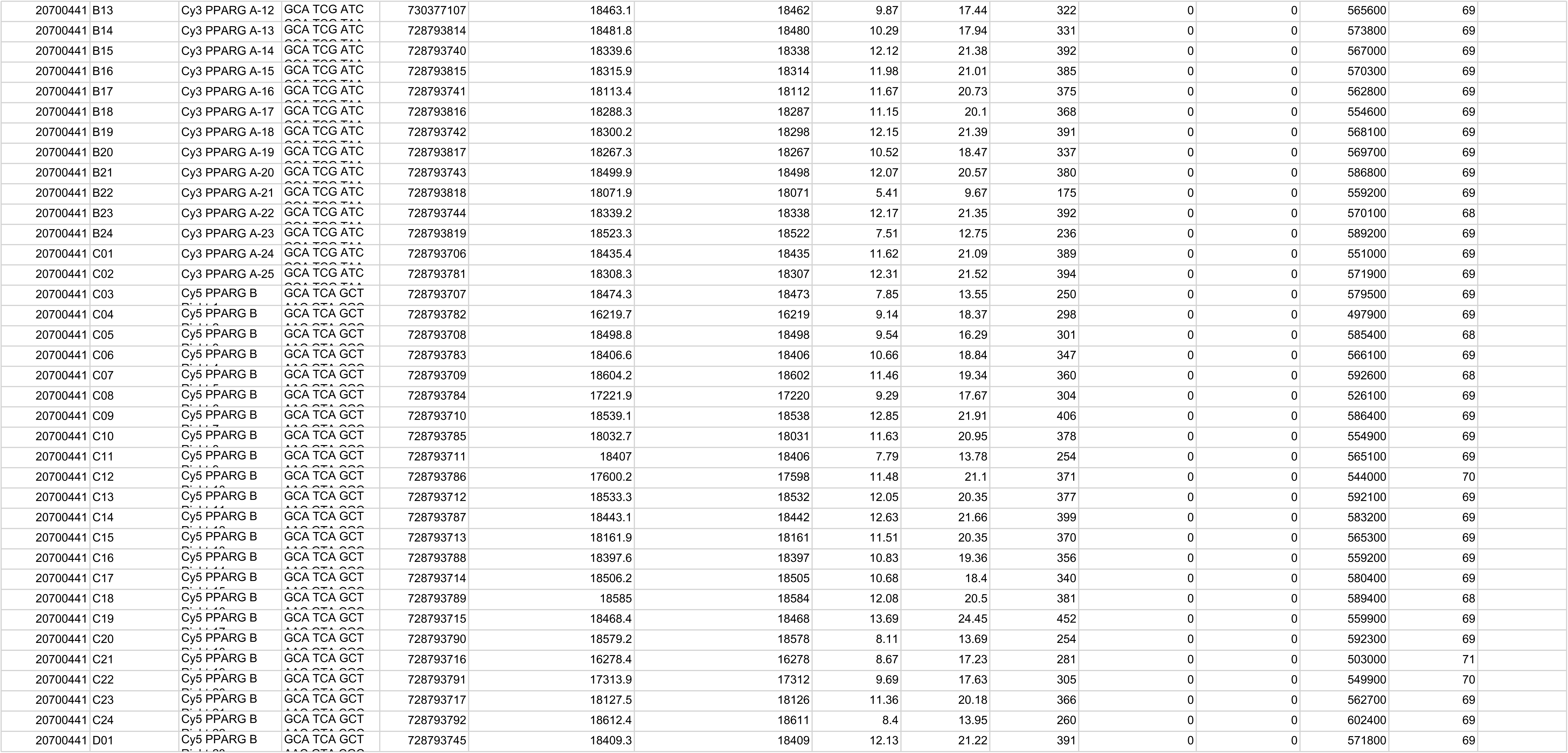
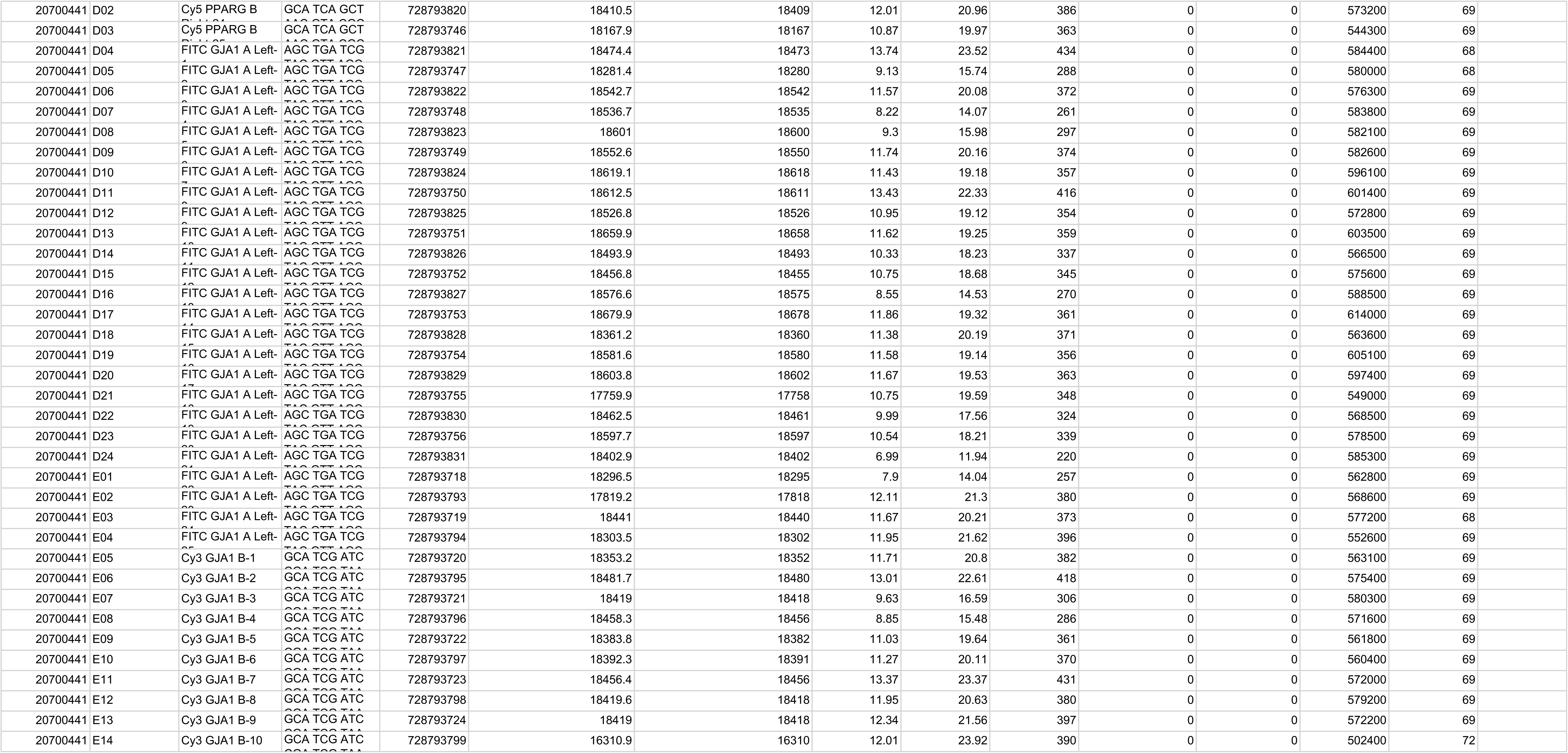
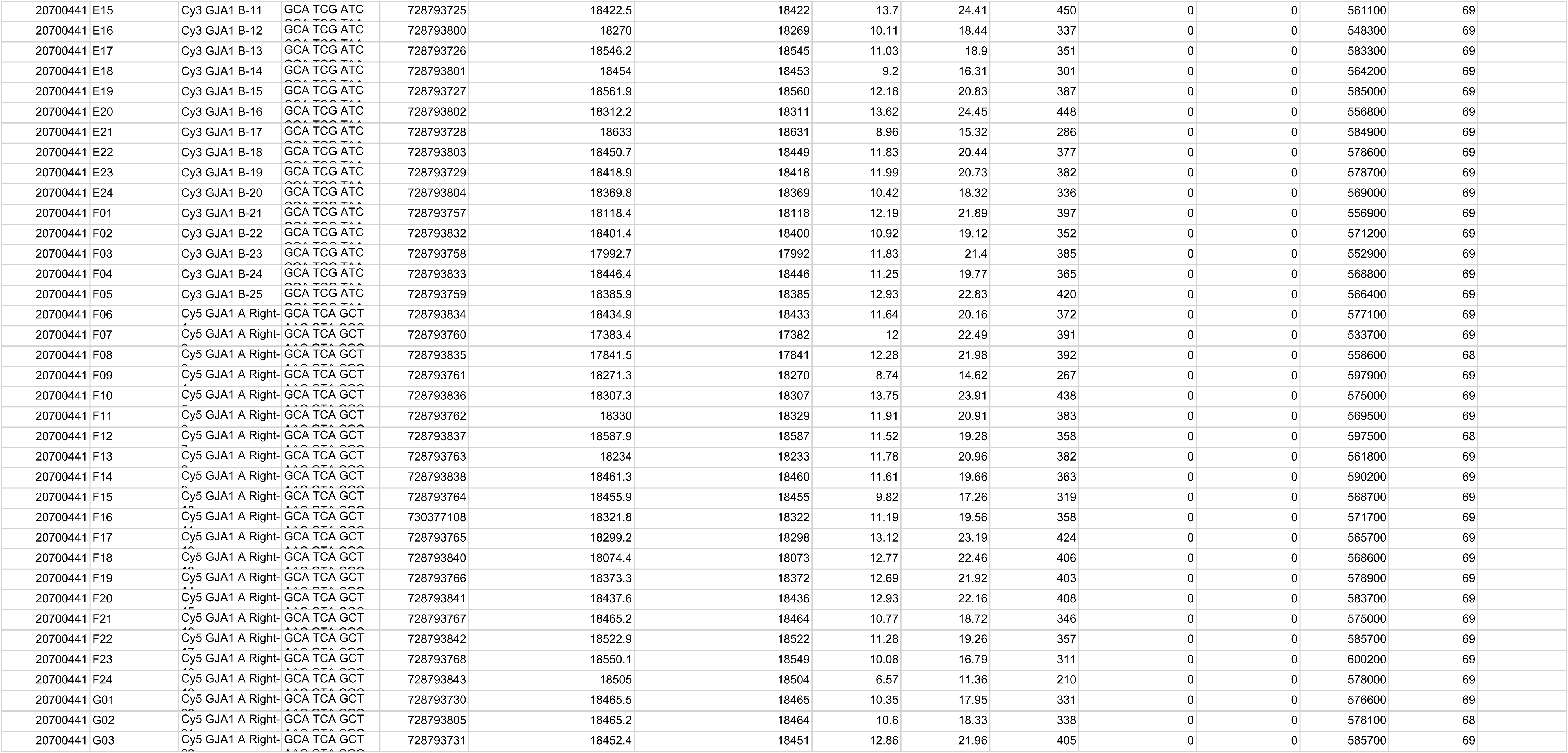
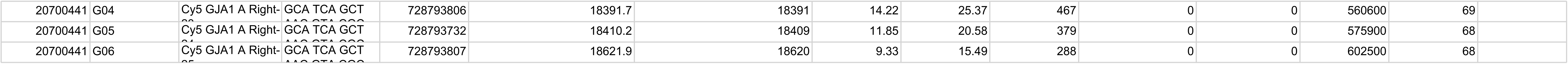

